# Cell-type specific effects of genetic variation on chromatin accessibility during human neuronal differentiation

**DOI:** 10.1101/2020.01.13.904862

**Authors:** Dan Liang, Angela L. Elwell, Nil Aygün, Michael J. Lafferty, Oleh Krupa, Kerry E. Cheek, Kenan P. Courtney, Marianna Yusupova, Melanie E. Garrett, Allison Ashley-Koch, Gregory E. Crawford, Michael I. Love, Luis de la Torre-Ubieta, Daniel H. Geschwind, Jason L. Stein

## Abstract

Common genetic risk for neuropsychiatric disorders is enriched in regulatory elements active during cortical neurogenesis. However, the mechanisms mediating the effects of genetic variants on gene regulation are poorly understood. To determine the functional impact of common genetic variation on the non-coding genome longitudinally during human cortical development, we performed a chromatin accessibility quantitative trait loci (caQTL) analysis in neural progenitor cells and their differentiated neuronal progeny from 92 donors. We identified 8,111 caQTLs in progenitors and 3,676 caQTLs in neurons, with highly temporal, cell-type specific effects. A subset (∼20%) of caQTLs were also associated with changes in gene expression. Motif-disrupting alleles of transcriptional activators generally led to decreases in chromatin accessibility, whereas motif-disrupting alleles of repressors led to increases in chromatin accessibility. By integrating cell-type specific caQTLs and brain-relevant genome-wide association data, we were able to fine-map loci and identify regulatory mechanisms underlying non-coding neuropsychiatric disorder risk variants.

**Highlights:**

- Genetic variation alters chromatin architecture during human cortical development
- Genetic effects on chromatin accessibility are highly cell-type specific
- Alleles disrupting TF motifs generally decrease accessibility, except for repressors
- caQTLs facilitate fine-mapping and inference of regulatory mechanisms of GWAS loci

## Introduction

Genome-wide association studies (GWAS) have revealed many common single nucleotide polymorphisms (SNPs) that are associated with risk for neuropsychiatric disorders (Cross-Disorder Group of the Psychiatric Genomics Consortium, 2013; Geschwind and Flint, 2015; Sullivan et al., 2018). A crucial next step is to understand the molecular mechanisms by which these variants impact disease risk (Sullivan and Geschwind, 2019). This is complicated by many factors, including: (1) risk loci are often comprised of many SNPs in high linkage disequilibrium (LD) and thus the causal variant(s) are not known (Barešić et al., 2019), (2) most risk loci have been mapped to non-coding regions with unknown, but presumed gene regulatory function (Lee et al., 2018b), (3) it is not known in which cell-type(s), tissue-type(s), or developmental time period(s) that a genetic risk variant exerts its effects (Gamazon et al., 2018). Nevertheless, a commonly assumed model to explain molecular mechanisms underlying risk loci is that non-coding risk alleles disrupt transcription factor (TF) binding motifs within cell-type specific regulatory elements (REs) leading to alterations in gene expression and downstream impacts on risk (Albert and Kruglyak, 2015; Nord and West, 2019). Thus, understanding tissue- and cell-type specific regulatory mechanisms are essential for moving from genetic association to a meaningful biological understanding.

With this in mind, several consortia including ENCODE, GTEx and PsychENCODE have taken major steps to build maps of non-coding genome function across the body (Davis et al., 2018; ENCODE Project Consortium, 2012; GTEx Consortium et al., 2017; Wang et al., 2018). These and other efforts have connected non-coding genetic variation to genes they regulate in developing and adult brain tissue by profiling 3-dimensional chromatin interactions and by measuring the impact of genetic variation on gene expression, termed expression quantitative trait loci (eQTLs) (Walker et al., 2019; Wang et al., 2018; Won et al., 2016). Although these studies are an important first step in connecting non-coding risk loci to their cognate genes, they do not elucidate the mechanisms by which human genetic variation alters gene regulation. Moreover, most of these resources focus on bulk tissue rather than specific cell-types, in which genetic variation can exert differing effects (Cuomo et al., 2019; Strober et al., 2019).

Risk variants for multiple neuropsychiatric disorders are enriched in REs active at mid-gestation in humans, during the peak of cortical neurogenesis (de la Torre-Ubieta et al., 2018). Histone acetylation QTL (haQTLs) and chromatin accessibility QTL (caQTLs) are powerful tools employed to identify the effect of genetic variation on non-coding REs and provide further understanding of gene regulatory mechanisms at both bulk and cell-type specific levels (Bryois et al., 2018; Degner et al., 2012; Gate et al., 2018; Schwartzentruber et al., 2018; Tehranchi et al., 2019, 2016; Wang et al., 2018). However, the ability to connect human allelic variation to longitudinal changes in regulatory architecture that occur during this foundational stage of human brain development is severely limited by the inaccessibility of brain tissue from the same individual over multiple time points. Here, we leveraged a well validated model of *in vivo* human brain development based on the *in vitro* culture of primary human neural progenitors (Dolmetsch and Geschwind, 2011; Stein et al., 2014) to study the functional effects of allelic variation on chromatin architecture during neurogenesis. We measured chromatin accessibility (Buenrostro et al., 2013) in neural progenitor cultures in a cell-type specific manner at two key stages of neural development, during progenitor proliferation (N_donor_= 73) and after differentiation using their labeled and sorted neuronal progeny (N_donor_= 61). We identified thousands of caQTLs, many of which were highly cell-type specific. We use the effects of these inherent genetic variations to understand how disrupting TF binding motifs impact chromatin accessibility and gene expression, as well as to understand the cell-type specific regulatory mechanisms underlying risk for neuropsychiatric disorders.

## Results

### Profiling genome-wide chromatin accessibility in progenitors and neurons

We generated primary human neural progenitor cell (phNPC) lines from 14-21 gestation week genotyped human fetal brain (N=92) using a neurosphere isolation method that results in cultures with high fidelity to the *in vivo* developing brain (Konopka and Wexler, 2010; Palmer et al., 2001; Rosen et al., 2011; Stein et al., 2014; Wexler et al., 2011) (Figure 1A; Methods). phNPCs were cultured and isolated at two stages for downstream experiments: progenitor cells and 8-week differentiated and sorted neurons. Over 90% of the undifferentiated progenitor cells were positive for SOX2 and PAX6 via immunofluorescence, indicating a highly homogenous population of radial glia cells (Figure 1A) (Hansen et al., 2010). After differentiation for 8 weeks, we FACS sorted the virally labeled differentiated neurons which showed typical neuronal morphology (Figure 1A; Supplemental Figure 1A; Methods). We performed ATAC-seq on intact nuclei and found that resultant libraries were high quality based on a sensitivity analysis and nucleosome periodicity (Supplemental Figure 1B-1D; (Buenrostro et al., 2013)). We quantified accessibility as batch effect-corrected reads within accessible peaks normalized for GC content, peak length and sequence depth (Supplemental Figure 2A and 2C-2D). We found higher correlations of chromatin accessibility from libraries prepared from the same donors cultured at different times as compared to correlations across donors (Progenitor: average within-donor correlation was 0.94 (± s.d. 0.009), average across-donor correlation was 0.91 (± s.d. 0.03); Neuron: average within-donor correlation was 0.93 (± s.d. 0.03), average across-donor correlation was 0.90 (± s.d. 0.04); Supplemental Figure 2B). To ensure independence for subsequent analyses, we randomly selected one library from each donor for each cell-type (N_progenitor=86 and N_neuron=67) to identify accessible peaks (N=136,714; average peak length of 508 bp; Methods).

**Figure 1:**
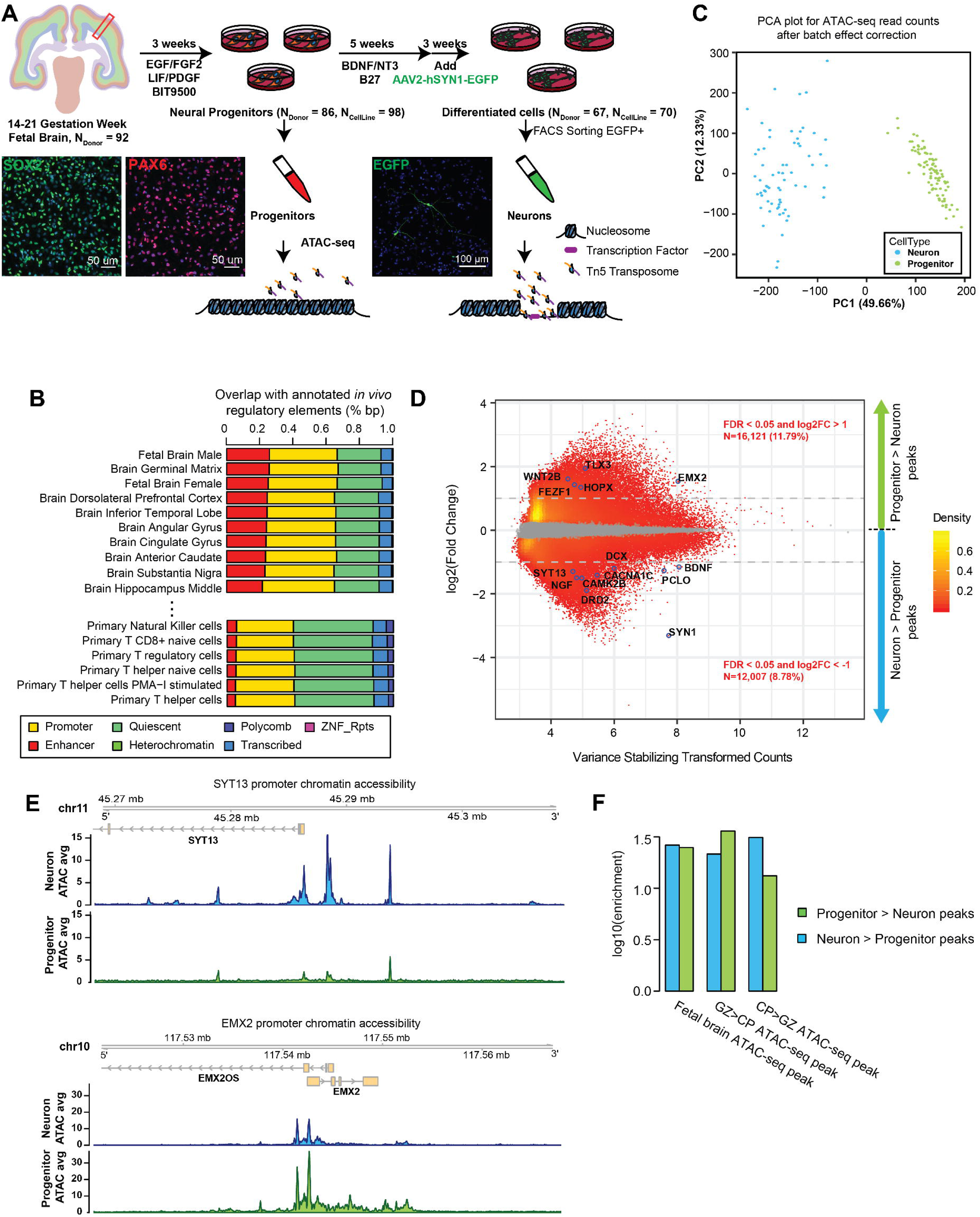
Profiling genome-wide chromatin accessibility in progenitors and neurons. (A) Schematic cartoon of experimental design. SOX2 and PAX6 immunolabeled neural progenitors (left), showing over 90% of cells were radial glia. EGFP labeled differentiated neurons (right), showing expected neuronal morphology. (B) Percentage of accessible peak base pairs (bp) detected in these cultures overlapped with with epigenetically annotated regulatory elements from multiple tissues. From top to bottom, tissues ordered by bp percentage overlap with enhancers. (C) PCA plot of ATAC-seq read count after batch effect correction colored by cell types, showing two separate clusters for progenitors and neurons. (D) MA plot for differentially accessible peaks between progenitors and neurons. All peaks can be found in Table S1. (E) ATAC-seq coverage plot (average normalized read counts) for promoter of neuron expressed gene SYT13, showing higher accessibility in neurons than progenitors (LFC=-1.29, FDR=3.87e-33). ATAC-seq coverage plot (average normalized read counts) for promoter of progenitor expressed gene EMX2, showing higher accessibility in progenitors than neurons (LFC=1.53, FDR=4.37e-24). (F) Enrichment of differentially accessible peaks identified here with differentially accessible peaks from fetal brain tissue (de la Torre-Ubieta et al., 2018). GZ: neural progenitor-enriched region encompassing the ventricular zone (VZ), subventricular zone (SVZ), and intermediate zone (IZ); CP: the neuron-enriched region containing the subplate (SP), cortical plate (CP), and marginal zone (MZ).

To determine the *in vivo* relevance of these accessible peaks, we performed an overlap analysis with REs derived from different human tissues (Figure 1B), utilizing previously classified chromatin states from 93 *in vivo* tissues and cell types (Roadmap Epigenomics Consortium et al., 2015), including fetal brain (male and female). The accessible peaks generated here from progenitors and neurons most strongly overlap enhancers identified in fetal brain tissue (Rank of enrichment across tissues: 1st for male (25.7% overlap) and 3rd (25.0% overlap) for female fetal brain), followed by other brain regions such as germinal matrix and dorsolateral prefrontal cortex, indicating that these peaks derived from phNPC cultures were highly representative of the *in vivo* fetal brain. The principal component analysis of chromatin accessibility revealed that progenitor and neuron samples clearly separate along the first principal component (Figure 1C), indicating that cell-type was associated with the largest variability in global chromatin accessibility profiles (49.66% of variance explained). These results demonstrate that chromatin accessibility measured from hNPC cultures are representative of REs present in the developing human brain and that chromatin accessibility patterns are broadly different between progenitors and neurons, consistent with previous bulk tissue data from fetal brain (de la Torre-Ubieta et al., 2018).

### Identifying cell-type specific regulatory elements during neuronal differentiation

To reveal cell-type specific REs involved in neuronal differentiation, we performed an analysis to determine which peaks had significantly different chromatin accessibility between neural progenitors and neurons (Figure 1D; Methods). We identified 16,121 peaks with greater accessibility in progenitors than neurons (progenitor peaks; FDR < 0.05 and log2 fold change (LFC) > 1) and 12,007 peaks with greater accessibility in neurons than progenitors (neuron peaks; FDR < 0.05 and LFC < −1; Table S1). At the promoter of *SYN1*, which was used to label neurons for sorting during differentiation, we observed considerably higher accessibility in neurons as compared to progenitors, as expected (LFC=-3.32, P-value=2.71e-59; Figure 1D). Among significant differentially accessible peaks, we found greater accessibility in progenitors at the promoters of genes highly or uniquely expressed in progenitors, including *WNT2B*, *NOTCH1*, *HOPX* and the dorsal telencephalic marker gene *EMX2* (Figure 1E) (Brunelli et al., 1996; Gaiano et al., 2000; Harrison-Uy and Pleasure, 2012; Pollen et al., 2015; Simeone et al., 1992). Moreover, promoters of genes highly expressed in neurons, such as *DCX*, *BDNF*, *CAMK2B* and *SYT13* (Fukuda and Mikoshiba, 2001; Polioudakis et al., 2019), showed greater chromatin accessibility in neurons (Figure 1D,E).

We identified the biological processes involved in differentially accessible peaks near protein-coding TSSs (2kb upstream and 1kb downstream from TSSs) during neuronal differentiation using Gene Ontology (GO) terms ((Ashburner et al., 2000); Supplemental Figure 3A). In promoter peaks more accessible in progenitors, we found an enrichment of GO terms related to known progenitor cell function, including forebrain development and neuron projection guidance. Conversely, in promoter peaks more accessible in neurons, we found enrichment of GO terms related to synaptic function, including signal release from synapse and neurotransmitter secretion, as well as learning, consistent with biological processes central to neuronal function. These results demonstrated that chromatin accessibility differences correspond to expected cell-type specific biological processes.

We also found that differentially accessible peaks were significantly enriched in ATAC-seq peaks from the relevant *in vivo* fetal brain laminae (Figure 1F) (de la Torre-Ubieta et al., 2018). Specifically, peaks more accessible in progenitors were more enriched in peaks with higher accessibility in the neural progenitor-enriched germinal zone. Conversely, peaks more accessible in neurons were more enriched in peaks more accessible in the neuron-enriched cortical plate. These comparisons showed the differentially accessible peaks represent cell-type specific active REs and were in strong agreement with biological processes and gene regulatory behavior present in *in vivo* fetal brain tissues.

To detect TFs involved in neuronal differentiation, we conducted a differential motif enrichment analysis to identify predicted TF binding sites more active in either progenitors or neurons. We detected 62 TFs (FDR < 0.05) with binding sites present more often in progenitor peaks than neuron peaks (here, called progenitorTFs), and 241 TF motifs presents more often in neuron peaks than progenitor peaks (here, called neuronTFs) (Methods; Supplemental Figure 3B). Within progenitorTFs, we found TFs previously characterized to have key roles for neural stem cell renewal and reprogramming, such as *SOX2* and *KLF4* (Ellis et al., 2004; Han et al., 2012; Wang et al., 2015), and those known to be required for the maintenance of stem cells in cortex, such as *NR2F1*, *ETV5*, and *SP2* (Liang et al., 2013; Liu and Zhang, 2019; Naka et al., 2008). Within neuronTFs, *NEUROG2* and *LMX1A* were identified, which are known to drive neuronal differentiation (Araújo et al., 2018; Fathi et al., 2015), as well as TFs shown to induce neuronal identity from fibroblasts, including *ASCL2* and the *POU* family (Tsunemoto et al., 2018). NeuronTFs also included *CUX2*, a marker for layer II-III neurons (Cubelos et al., 2015; Zimmer et al., 2004) and other laminar markers such as *TBR1* and *FOXP1/2*. Thus, motifs within differentially accessible peaks allow the identification of known TFs that are essential for neural progenitor proliferation and differentiation *in vivo*, providing further support that TF binding within chromatin accessibility peaks in this *in vitro* system reflect the expected *in vivo* developmental processes (Supplemental Figure 3C). We also note that we identified several TFs that have not been previously associated with neuronal differentiation, such as *MEF2A*, *MIX-A* and *HOXD8*, which may be useful for directed differentiation of human neural progenitor cells (Table S2).

### Chromatin accessibility quantitative trait loci (caQTLs)

To identify genetic variants that influence chromatin accessibility within two key cell types present during cortical development, we performed caQTL analyses separately for progenitors and neurons (Figure 2A; Supplemental Figure 4A-4B). Because our sample was composed of multiple ancestries (Supplemental Figure 4C), we stringently controlled for population stratification in our association tests using a linear mixed effects model including a kinship matrix as a random effect, and 10 genotype MDS components as fixed effects (Kang et al., 2010; Price et al., 2010; Yang et al., 2014). In addition, we included sex, gestation week, and principal components (PCs) across the chromatin accessibility matrix (4 PCs in neurons; 7 PCs in progenitors) as fixed effect covariates to reduce the impact of unmeasured technical variation (Pickrell et al., 2010).

**Figure 2:**
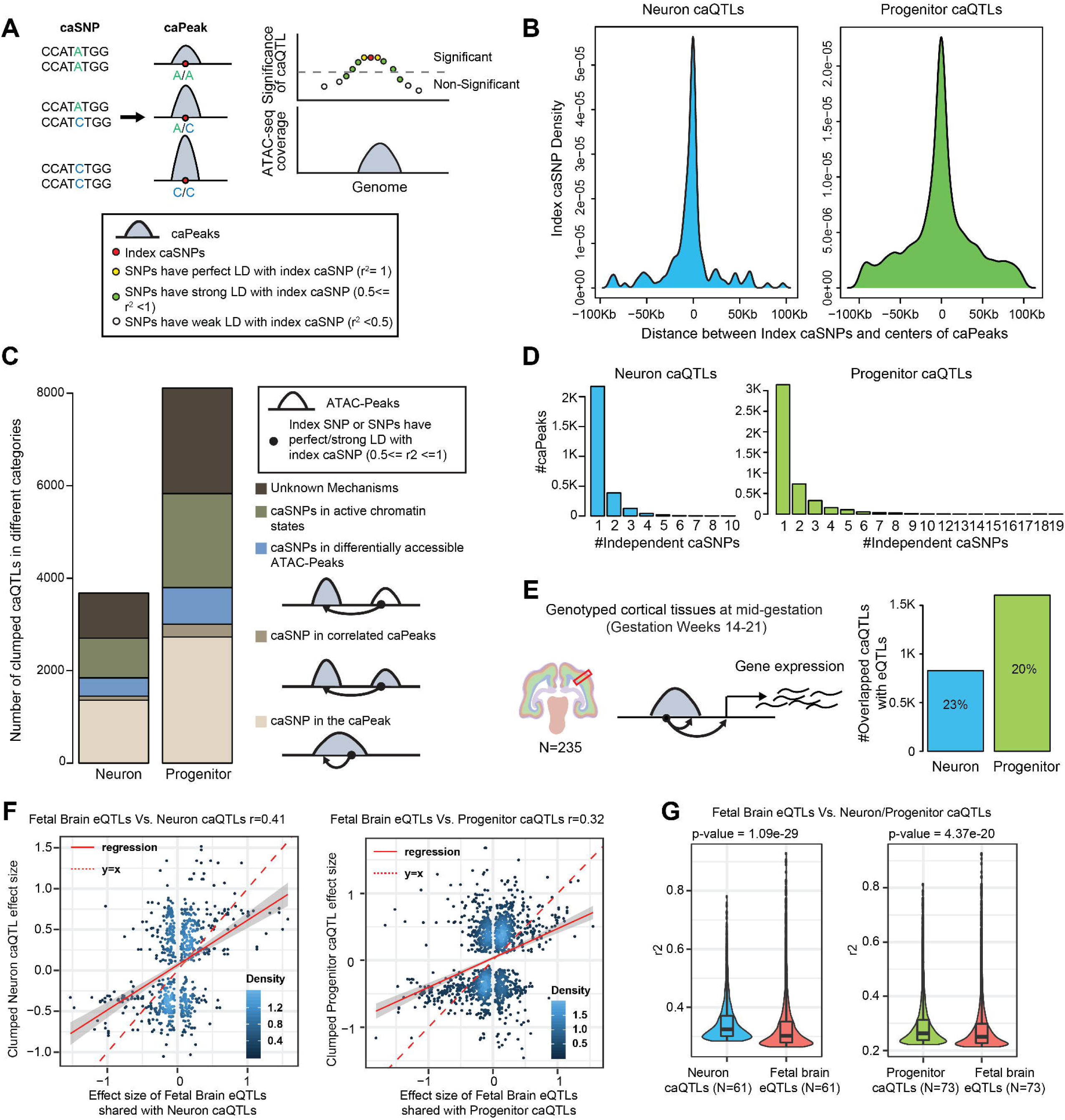
Chromatin accessibility quantitative trait loci (caQTL) in progenitors and neurons. (A) caQTL schematic. (B) Density of index caSNPs relative to the distance from the center of the caPeaks (left: neuron caQTLs; right: progenitor caQTLs). All clumped caQTLs can be found in Table S3. (C) Numbers of clumped caQTLs in different functional categories. (D) Numbers of independent caSNPs affecting chromatin accessibility of a caPeak. (E) Schematic cartoon of fetal cortical eQTLs (Walker et al., 2019) (Left). Numbers and percentage of clumped overlapped neuron/progenitor caQTLs with fetal cortical eQTLs. All overlapped ca/eQTLs can be found in Table S4. (F) Correlation of effect sizes between overlapped fetal cortical eQTLs and neuron (*left*) and progenitor (*right*) caQTLs. (G) Comparison of percent variance explained (r^2^) for caQTLs and eQTLs (subset to the same sample size) in neurons and progenitors. We observed higher percent variance explained for caQTLs than eQTLs in both neurons and progenitors.

We identified 8,111 LD-independent caQTLs (caSNP-caPeak pairs) regulating 4,682 caPeaks in progenitors (N_donor_=73) and 3,676 LD-independent caQTLs regulating 2,760 caPeaks in neurons (N_donor_=61). We found that the caSNP or an LD-proxy was located in a known functional region for ∼73% of caQTLs detected (Figure 2C; Table S3). These caPeaks were significantly enriched in active REs such as enhancers and promoters from fetal brain (Supplemental Figure 4D), consistent with their expected regulatory function. Index caSNPs are most often found near the peaks they are associated with (Figure 2B), half of index caSNPs of progenitor caQTLs are located within −24.3kb to 19.1kb around peak centers. Similarly, half of index caSNPs of neuron caQTLs are located within −27.5kb to 20.2kb around peak centers (Figure 2B). We found that one caPeak could be affected by multiple genetic variants (Figure 2D), indicating that caSNPs could disrupt multiple TF motifs within one regulatory element or the caPeak was regulated by other REs, including correlated peaks, differentially accessible peaks, or active chromatin regions. These results imply that most genetic variants affect chromatin accessibility by altering the sequence (and presumably transcription factor binding sites) at the peak impacted by the variant or disrupt chromatin accessibility at distal peaks which have secondary effects on the caPeak (Kumasaka et al., 2019).

To identify if genetic influences on chromatin accessibility were also associated with differences in gene expression, we compared progenitor and neuron caQTLs with recent eQTLs from the mid-gestation cortical wall (Walker et al., 2019) (Methods). Twenty percent of progenitor caQTLs and 23% of neuron caQTLs were shared with fetal cortical eQTLs (Figure 2E, Table S4). The effect sizes of shared caQTL-eQTL pairs showed positive correlations (r=0.41 for fetal cortical wall eQTLs and neuron caQTLs; r=0.32 for fetal cortical wall eQTLs and progenitor caQTLs). Among shared caQTL-eQTL pairs, alleles have the same direction of effect on chromatin accessibility and gene expression for most shared caQTL-eQTL pairs (63.4% of shared progenitor caQTLs and 64.5% shared neuron caQTLs), indicating that alleles associated with increased chromatin accessibility tend to be associated with increased gene expression (Figure 2F). For 36.6% of shared progenitor caQTLs and 35.5% shared neuron caQTLs, alleles have the opposite direction of effect on chromatin accessibility and gene expression, indicating a repressive function of these caPeaks on eGenes. We subsampled the eQTL dataset to ensure that caQTLs and eQTLs have the same sample sizes (Methods), and then assessed whether genetic variation explained more variation in chromatin accessibility as compared to gene expression (Figure 2G), observing that caQTLs generally explain more variance than eQTLs (neurons: average caQTL r^2^=0.350 and average eQTL r^2^=0.332, t-test p-value=1.088e-29; progenitors: average caQTL r^2^=0.292 and average eQTL r^2^=0.281, t-test p-value=4.369e-20). This indicates that caQTL studies may have higher power than eQTL studies and/or that cell-type specificity increases power for molecular-level genetic associations (Gate et al., 2018).

### Genetic effects on cell-type specific regulatory elements

By integrating cell-type specific caQTLs and fetal cortical eQTLs, we were able to annotate cell-type specific REs active during human neuronal differentiation and identify their cognate genes. We subsequently leveraged these data to fine map causal variants within eQTL loci and predict regulatory mechanisms of these non-coding SNPs associated with both chromatin accessibility and gene expression. We identified 161 RE-Gene pairs in neurons and 285 RE-Gene pairs in progenitors. Within these RE-Gene pairs, we found many genes involved in neuronal differentiation such as *SCN1A, FABP7, NEK3* and *VAT1* (Chang et al., 2009; Escayg and Goldin, 2010; Feng et al., 1994; Loeb-Hennard et al., 2004). We also identified RE-gene pairs where the caSNP disrupted motifs of TFs that have known functions in early stem cell differentiation and neuronal differentiation (Ballas et al., 2005; Nitzsche et al., 2011). For example, the C allele of rs11544037 is associated with increased chromatin accessibility of a progenitor caPeak (chr4:158,667,741-158,667,880) located ∼5 kb upstream from the *ETFDH* TSS and also associated with increased expression of *ETFDH* (Figure 3A-3C). We found that rs11544037 disrupted several TF motifs. These TFs were ranked by their level of expression in progenitors and neurons using fetal brain scRNA-seq data (Polioudakis et al., 2019). We prioritized RAD21 as the TF with altered binding due to the caSNP based on its high level of expression in progenitors (Figure 3D). The C allele of rs11544037 matched the motif of RAD21 (Figure 3E), which suggests that RAD21 binding was associated with increased chromatin accessibility of this enhancer in progenitors and increased expression of *EFTDH*. Another example was the G allele of rs185220 which was associated with increased chromatin accessibility in neurons and progenitors of a caPeak (chr5:56,909,071-56,910,960) that overlapped with the *SETD9* TSS and was associated with increased expression of *SETD9* (Figure 3F-3H). We found that rs185220 disrupted several TF motifs, but we prioritized *REST* due to its high expression in progenitors ((Polioudakis et al., 2019) Figure 3I-J). In contrast to the previous example, the G allele of rs185220 led to disruption of the REST motif and increased chromatin accessibility, consistent with the function of REST as a repressor (Chong et al., 1995; Schoenherr and Anderson, 1995). The predicted regulatory mechanism of rs185220 is that the G allele of rs185220 disrupted REST binding, resulting in increased chromatin accessibility and increased expression of *SETD9*.

**Figure 3:**
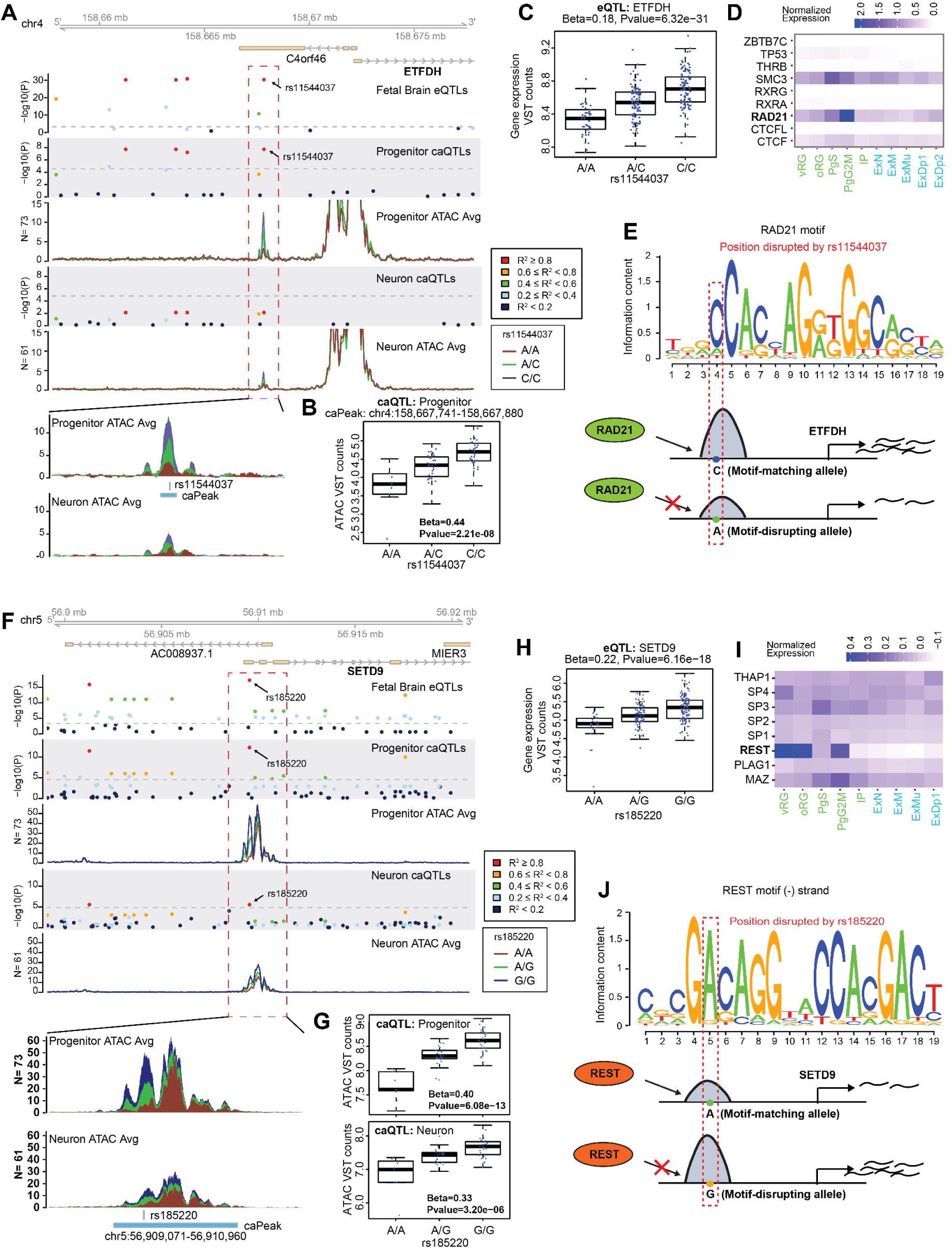
Fine-mapping and regulatory mechanism underlying eQTLs. (A) Co-localization of a progenitor-specific caQTL and the fetal cortical eQTL for *ETFDH*. (B) The association between rs11544037 and chromatin accessibility of the labeled peak. (C) The association between rs11544037 and expression of *ETFDH* in fetal cortex. (D) The expression of TFs in which motifs are disrupted by rs11544037. vRG: ventricular Radial Glia; oRG: outer Radial Glia; PgS: Cycling progenitors (S phase); PgG2M: Cycling progenitors (G2/M phase); IP: Intermediate progenitors; ExN: Migrating excitatory; ExM: Maturing excitatory; ExM-U: Maturing excitatory upper enriched; ExDp1: Excitatory deep layer 1; ExDp2: Excitatory deep layer 2. (E) The motif Logo of *RAD21*, where the red box shows the position disrupted by rs11544037. Schematic cartoon of mechanisms for rs11544037 regulating chromatin accessibility and gene expression. (F) Co-localization of a caQTL in progenitors and neurons and the fetal cortical eQTL for *SETD9*. (G) The association between rs185220 and chromatin accessibility of the labeled peak in progenitors (top) and neurons (bottom). (H) The association between rs185220 and expression of *SETD9* in fetal cortex. (I) The expression of TFs in which motifs are disrupted by rs185220. (J) The motif Logo of *REST*, where the red box shows the position disrupted by rs185220. Schematic cartoon of mechanisms for rs185220 regulating chromatin accessibility and gene expression.

### Allele Specific Chromatin Accessibility

We next performed an analysis to detect allele specific chromatin accessibility (ASCA) for each heterozygous SNP located within accessible peaks (Methods). ASCA analysis contrasts accessibility between two alleles within an individual heterozygous at a given SNP, so it inherently controls for cross-individual confounding factors, such as population structure (Pastinen, 2010). In total, we identified 1,598 significant (FDR < 0.05) progenitor ASCA and 3,332 significant neuron ASCA (Table S5). To determine if caQTLs also show ASCA, we filtered to keep significant caQTLs (non-clumped, FDR < 0.05) using the same heterozygous donor and read level criteria described for ASCA, observing that 91.2% of filtered neuron caQTLs were shared with neuron ASCA (Fisher’s test: OR=56.76, p-value=3.9e-207) and 87.7% of filtered progenitor caQTLs were shared with progenitor ASCA (Fisher’s test: OR=48.97, p-value=1.6e-215). This demonstrates extremely high overlap between caQTLs and ASCA (Figure 4A), which indicates minimal influence of cross-individual confounding effects, such as population stratification, on the caQTL results. Similarly, for all filtered caQTLs and significant ASCA in Figure 4A, we found high correlations of effect sizes between caQTLs and ASCA (r=0.67 for neurons; r=0.75 for progenitors), indicating a shared direction and degree of effect (Figure 4B). Subsetting to only significant caQTLs and ASCAs, the correlation of effect sizes is greater than 0.9 for both neurons and progenitors (Figure 4B), again consistent with the lack of confounding due to population factors for most caQTLs discovered.

**Figure 4:**
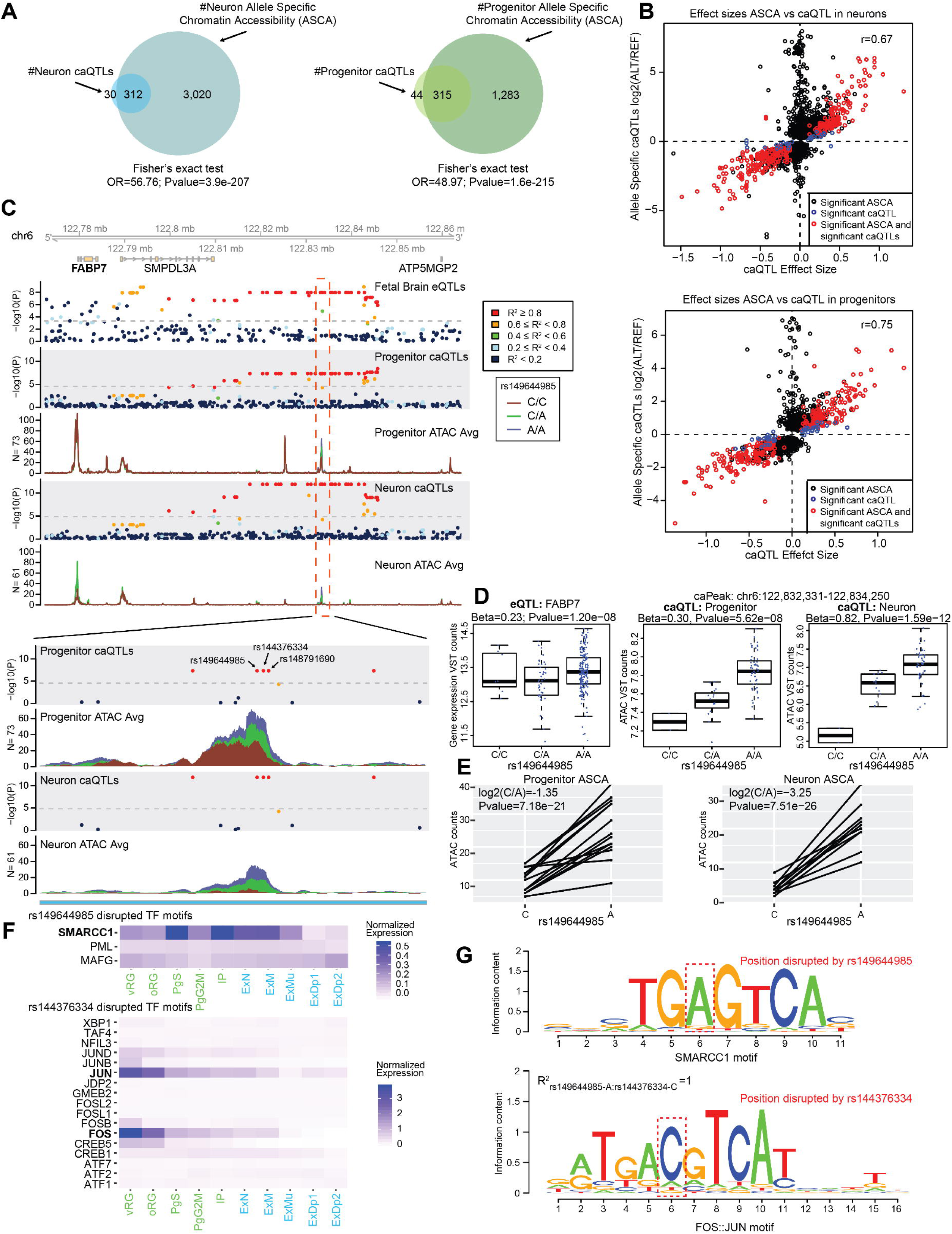
Allele Specific Chromatin Accessibility (ASCA). (A) Numbers of shared/non-shared significant caQTLs and significant ASCA in neurons (left) and progenitors (right). All significant ASCA in neurons and progenitors can be found in Table S5. (B) Correlation of effect sizes for caQTL and ASCA from (A) in neurons (*top*) and progenitors (*bottom*). (C) Co-localization of caQTL and ASCA in progenitors and neurons as well as fetal cortical eQTL for *FABP7*. (D) Association between rs149644985 and expression of *FABP7* in fetal cortex (*left*), chromatin accessibility of the labeled peak in progenitors (*middle*) and neurons (*right*). (E) ASCA detected at rs149644985 in progenitors (*left*) and neurons (*right*). (F) The expression of TFs where motifs are disrupted by rs149644985 (*top*). The expression of TFs where motifs are disrupted by rs144376334 (*bottom*). (G) The motif logo of SMARCC1 disrupted by rs149644985 (*top*); the motif logo of FOS::JUN disrupted by rs144376334 (*bottom*).

However, we also detected significant ASCAs that were not significant caQTLs (Figure 4B). These variants were found in larger peaks than those detected in both caQTL and ASCA (Supplemental Figure 5A). These ASCA-but-not-caQTL variants likely have an effect on the accessibility of a sub-region of the larger active region. They are more detectable using ASCA because only reads containing the variant at the location where accessibility is affected are tested for association, whereas they are not detectable in caQTLs which integrate reads across the entirety of the region. For example, we detected ASCA at SNP rs2547972, which was associated with differences in chromatin accessibility of a sub-peak, but was not a caQTL for the called larger peak where it resides (9,139 bp; Supplemental Figure 5B). Other ASCA, but not caQTL sites, were presumably due to lower power for caQTL detection (Supplemental Figure 5C).

We identified several loci that shared caQTLs, ASCA, and eQTLs. For example, the previously described *SETD9* locus also demonstrated strong ASCA at rs185220 in both neurons and progenitors (Supplemental Figure 5D). We were also interested in *FABP7* (also known as BLBP), which is a marker for radial glia that plays an important role in the establishment of the radial glial fibers spanning the cortical anlage during cortical development (Feng et al., 1994) (Figure 4C). The A allele of rs149644985 (and its LD proxies found within the same caPeak rs144376334 r^2^_rs149644985-A:rs144376334-C_=1 and rs148791690 r^2^_rs149644985-A:rs148791690-C_=1) was associated with increased chromatin accessibility of the caPeak (chr6:122,832,331-122,834,250) in both progenitors and neurons and increased gene expression of *FABP7* (Figure 4D). The A allele of rs149644985 (C allele of rs144376334) also manifested increased allele-specific chromatin accessibility in both progenitors and neurons (Figure 4E). rs149644985 and rs144376334 disrupted several TF motifs, and we prioritized SMARCC1 and FOS::JUN due to their higher expression in progenitors (Polioudakis et al., 2019) (Figure 4E). The motif disrupting allele for both SNPs was associated with decreased chromatin accessibility, consistent with activating REs (Figure 4G). Based on these data, we suggest two potential regulatory mechanisms underlying one genetic locus based on different disrupted TF motifs: 1) genetic variation simultaneously disrupts both SMARCC1 and FOS::JUN binding to the RE leading to decreased expression of FABP7 or 2) genetic variation disrupting either SMARCC1 or FOS::JUN is sufficient to decrease the expression of *FABP7*.

### Genetic effects on chromatin accessibility are cell-type specific

To determine the cell-type specificity of caQTLs between progenitors and neurons, we quantified the degree of overlap using LD-independent progenitor and neuron caQTLs. We identified if any neuron caSNPs (or their LD proxies; 0.5 <= r^2^ <=1) overlapped with progenitor caSNPs (or their LD proxies), and the converse. Only 10.8% of progenitor caQTLs (caSNP-caPeak pairs) overlapped with neuron caQTLs and 17.4% of neuron caQTLs overlapped with progenitor caQTLs (Figure 5A). To determine if genetic variation impacts the same peaks in both progenitors and neurons, we quantified the degree of overlap in caPeaks. We found only 8.9% of progenitor caPeaks overlapped with neuron caPeaks, and that 15.1% of neuron caPeaks overlapped with progenitor caPeaks (Figure 5B). For ASCA, we found 24.2% progenitor ASCA and 17.6% neuron ASCA are shared, which was in agreement with the cell-type specificity observed in caQTLs. These results suggest that genetic variants often impact chromatin accessibility only within specific cell-types.

**Figure 5:**
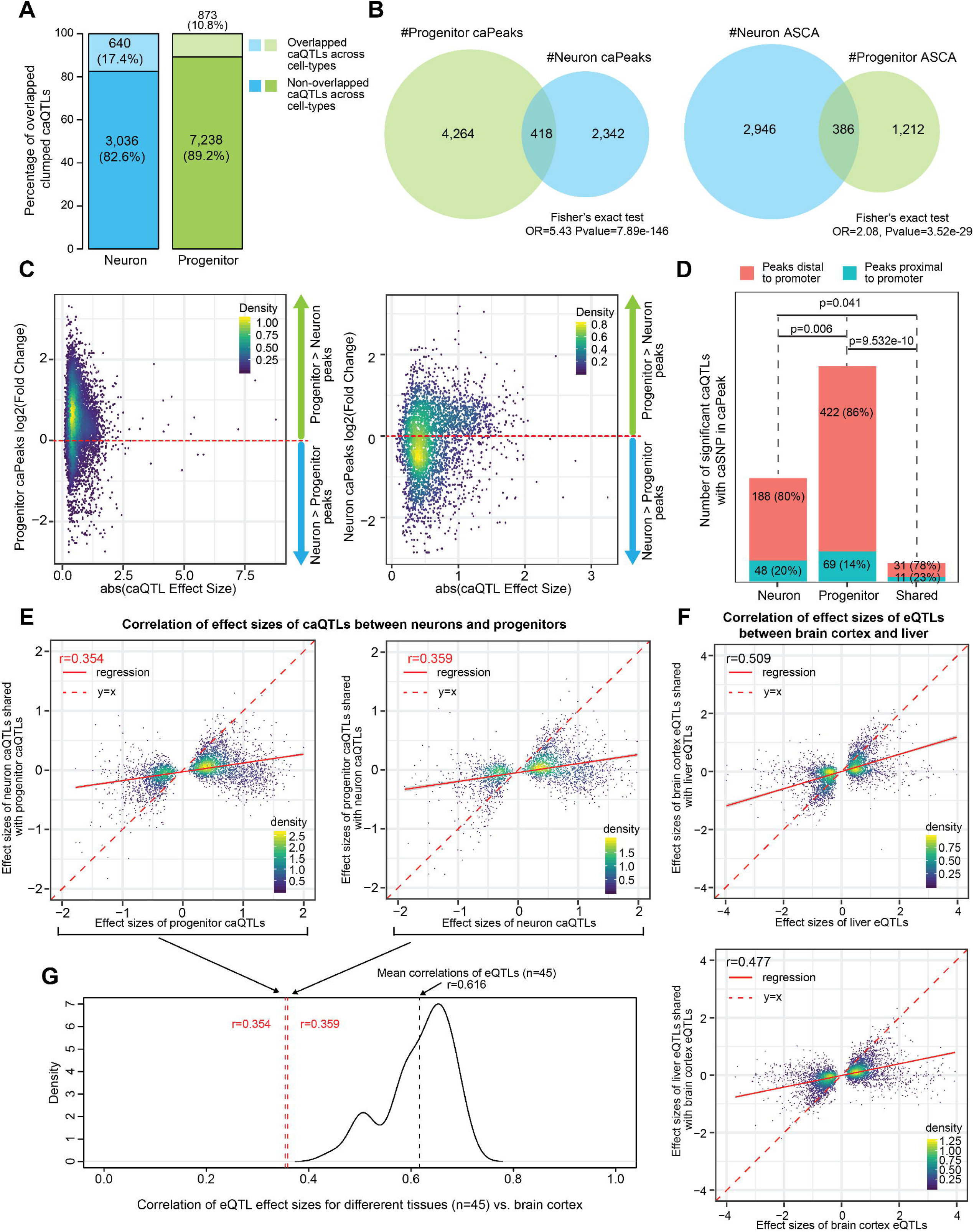
Cell-type specificity of caQTLs. (A) Numbers and percentage of overlapped caQTLs between neurons and progenitors. (B) Numbers of overlapped/non-overlapped caPeaks (*left*) and ASCA (*right*) between neurons and progenitors. (C) Differential accessibility of progenitor caPeaks (*left*) and neuron caPeaks (*right*). (D) Numbers and percentage of caPeaks distal to promoters or proximal to promoters for neuron-specific significant caQTLs, progenitor-specific significant caQTLs and shared caQTLs between neurons and progenitors. (E) Correlations of effect sizes of caQTLs between neurons and progenitors (left: selected the most significant neuron caQTLs for every caPeak vs. progenitor caQTLs; right: selected the most significant progenitor caQTLs for every caPeak vs. neuron caQTLs). (F) Correlations of effect sizes of eQTLs between brain cortex and liver. (left: selected the most significant liver eQTLs for every eGene vs. brain cortex eQTLs; right: selected the most significant brain cortex eQTLs for every eGene vs. liver eQTLs). (G) Density plot for correlations of eQTL effect sizes between different tissues and brain cortex. The mean of these correlations is 0.616 which is much larger than correlations of caQTLs effect sizes between progenitors and neurons.

To further characterize the cell-type specificity of caQTLs, we assessed the differential accessibility of progenitor and neuron caPeaks (Figure 5C). We found the caPeaks of 75.4% of progenitor caQTLs were more accessible in progenitors as compared to neurons (LFC > 0). Similarly, the caPeaks of 52.6% neuron caQTLs were more accessible in neurons as compared to progenitors (LFC < 0). This implies genetic variants that disrupted the binding of DNA-binding proteins, like transcription factors, affected chromatin accessibility of the REs.

We next characterized the location of caPeaks relative to the nearest promoter (2kb upstream and 1kb downstream from TSS) when the caSNP is located within the caPeak (Figure 5D). We found there was a higher percentage of cell-type specific caPeaks (86% of progenitor caPeaks and 80% of neuron caPeaks) that were distal to promoters than shared caPeaks (78%; Progenitor: p=9.532e-10 and Neuron: p=0.041). This result indicates cell-type specific caQTLs were more likely to affect the chromatin accessibility of distal REs, which is consistent with distal REs such as enhancers having higher cell-type specificity than promoters (Heinz et al., 2015; Roadmap Epigenomics Consortium et al., 2015).

To determine the direction and magnitude of the effect of a genetic variant on chromatin accessibility between cell types, we related the effect sizes from the most significant caQTLs for every caPeak detected in one cell-type to the other. Consistent with the observation that caQTLs significant in one cell-type are rarely significant in another, we noted a relatively low correlation between effect sizes from the most significant neuron caQTLs compared with those same SNP-peak pairs in progenitors (r=0.359; p-value=1.056e-82; Figure 5E). Similarly, effect sizes from the most significant progenitor caQTLs showed a relatively low correlation with the same SNP-peak pairs in neurons (r=0.354; p-value=4.159e-127).

Previous research has shown surprisingly high consistency of genetic effects on expression across many tissues in the body (GTEx Consortium et al., 2017). We therefore sought to determine if caQTLs have higher tissue/cell-type specificity than eQTLs. We compared the effect size correlations between cell-types from caQTL analyses to effect size correlations between tissue types from eQTL analyses derived from GTEx. We selected the most significant liver eQTL for each eGene and compared effect sizes for the same SNP-gene pairs to brain cortex, finding considerably higher correlation (r=0.509, p-value=4.333e-237; Figure 5F) than those observed from caQTLs. The converse analysis, selecting the most significant brain eQTLs and comparing effect sizes for the same SNP-gene pairs to liver similarly yielded a stronger correlation (r=0.477, p-value=1.340e-292) than those observed from caQTLs. To more globally compare tissue-type specificity of eQTLs to the cell-type specificity of caQTLs, we used the same method to calculate effect size correlation between independent significant eQTLs from other tissues (N_tissues_=45, excluding highly related tissues - see Methods). We found the average effect size correlation is 0.616 between different tissues in the body and brain cortex, which is much higher than the effect size correlations observed between neuron and progenitor caQTLs (Figure 5G). Together, these results suggested that caQTLs were highly cell-type specific between progenitors and neurons, in comparison with eQTLs which were more correlated between different tissues and brain cortex.

### Comparison to adult dorsolateral prefrontal cortex (DLPFC) caQTLs

Previous work identified common variant associations with chromatin accessibility in adult post-mortem DLPFC using a sample of 272 individuals (Bryois et al., 2018). We tested whether caQTLs present during neuronal differentiation of the cortex modeling a prenatal time period were also present in adult cortex (Supplemental Figure 6A-6C). Only 2.4% of DLPFC caQTLs overlapped with neuron caQTLs and 19.9% of neuron caQTLs overlapped with DLPFC caQTLs. Similarly, only 4.7% of DLPFC caQTLs overlapped with progenitor caQTLs and 17.5% of progenitor caQTLs overlapped with DLPFC caQTLs. In general, there are few variants that have similar impacts on chromatin accessibility in the cortex during fetal development, as compared to adulthood. The peaks impacted by genetic variation (caPeaks) were shared with adult cortex in only 17.5% of neuron caPeaks and 16.2% of progenitor caPeaks (Supplemental Figure 6D). For the subset of genetic variants that shared effects on chromatin accessibility between neuronal differentiation and in adult cortex, there were very strong correlations in the effect sizes (r=0.92 for DLPFC and neurons; r= 0.89 for DLPFC and progenitors) (Supplemental Figure 6E). Together, these results indicate that caQTLs have high temporal specificity, as well as cell-type specificity, but for the subset of shared caQTLs similar effects are observed in brain at different stages of development.

### Prediction of disrupted transcription factor (TF) binding due to genetic variation

One favored model of how genetic variation leads to changes in chromatin accessibility is that SNPs disrupt TF motifs, decreasing the probability of TF binding to chromatin, and resulting in decreased chromatin accessibility (Behera et al., 2018; Gate et al., 2018). To determine which TF motifs are disrupted by cell-type specific caSNPs, we mapped known TF motifs to the sequence surrounding the neuron-specific/progenitor-specific caSNPs and determined if an allele at the caSNP sufficiently decreases the relative entropy of TF binding using the position possibility matrix (PPM; Methods; Table S6; (Coetzee et al., 2015)). Most often, caSNPs only disrupted one TF motif (Figure 6A). However, given the ubiquitousness of motifs in the genome, some caSNPs can disrupt motifs of as many as 93 TFs (Figure 6A). Conversely, we found one motif could be disrupted by many genetic variants in different peaks (the *SOX9* motif was disrupted by 141 caSNPs). In progenitors, a motif often disrupted by caSNPs (accounting for the number of motifs disrupted by SNPs in accessible peaks) and expressed in progenitors was *SMARCC1* (Figure 6B; (Polioudakis et al., 2019)), a member of the BAF complex known to be involved in fate decisions during neurogenesis (Ronan et al., 2013). In neurons, we found the motif of *DLX3*, which is a critical TF for induced neuronal cell reprogramming, was the most often disrupted by caSNPs (Figure 6B) (Wapinski et al., 2017). These results suggest that the TFs whose motifs are disrupted by caSNPs are involved in neural proliferation and differentiation, indicating that the genetic variants that impact the activity of REs by disrupting the binding of TFs play functional roles during these biological processes.

**Figure 6:**
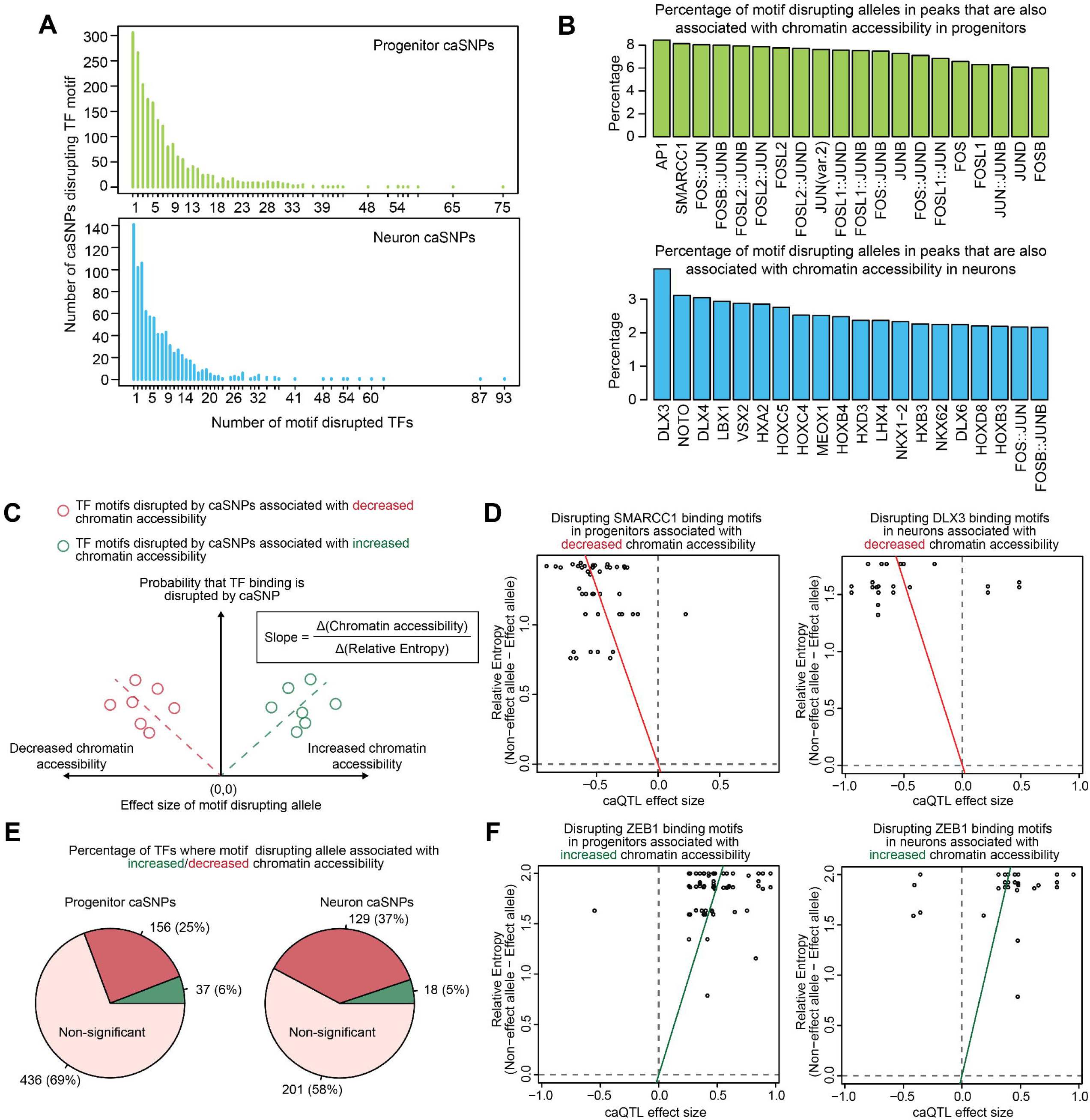
Prediction of disrupted transcription factor (TF) binding due to genetic variation. (A) Numbers of TFs where motifs were disrupted by progenitor-specific caSNPs (top) and neuron-specific caSNPs (bottom). (B) Percentage of motif disrupting alleles in accessible peaks that are also associated with chromatin accessibility in progenitors (*top*) or in neurons (*bottom*). (C) Schematic of TF motifs disrupted by caSNPs associated with decreased/increased chromatin accessibility. (D) Examples of TF motifs broken by caSNPs associated with decreased chromatin accessibility in progenitors (*SMARCC1*; left) and neurons (*DLX3*; right). (E) Numbers and percentage of TFs where the motif-disrupting allele was associated with increased/decreased chromatin accessibility in progenitors (*left*) and neurons (*right*). For most TFs, the motif-disrupting allele was associated with decreased chromatin accessibility in progenitors and neurons. (F) Disrupting *ZEB1* (a transcriptional repressor) binding motifs in progenitors was associated with increased chromatin accessibility in progenitors and neurons.

We next tested the impact of the TF motif-disrupting alleles on chromatin accessibility. We found alleles at caSNPs disrupting *SMARCC1* motifs are generally associated with decreased chromatin accessibility in progenitors (Methods; Figure 6C). Similarly, alleles at caSNPs disrupting *DLX3* motifs are associated with decreased chromatin accessibility in neurons (Figure 6D). We then performed this analysis using all available TF motifs where we observed at least 10 motif disrupting caSNPs. Among 629 tested TFs in progenitors and 348 tested TFs in neurons, we found that motif disrupting alleles often led to decreased accessibility for 156 (25%) TFs in progenitors and 129 (37%) TFs in neurons (Figure 6E). Conversely, we found the caSNP motif-disrupting allele was associated with increased chromatin accessibility at the motif of *ZEB1*, a known transcriptional repressor (Figure 6F) (Wang et al., 2019). Interestingly, we found the motif of *TCF12* in progenitors was disrupted by caSNPs associated with increased chromatin accessibility. However, in neurons, this motif was disrupted by caSNPs associated with decreased chromatin accessibility. *TCF12* is a basic helix loop helix (bHLH) transcription factor (Zhang et al., 1991) involved in the expansion of precursor cell populations during neurogenesis (Uittenbogaard and Chiaramello, 2002). *TCF12* forms homo and heteroduplex complexes with other bHLH factors and can act both as a repressor and activator of transcription depending on the cellular context (Hu et al., 1992; Wu et al., 2012; Zhang et al., 1991). Based on these results, we hypothesize that *TCF12* functions as a repressor of transcription in progenitors, but an activator in neurons. These results suggest that binding of transcriptional activators was associated with increased chromatin accessibility. However, binding of transcriptional repressors was associated with decreased chromatin accessibility.

### Regulatory mechanisms underlying GWAS loci

To investigate if genetic variants associated with common neuropsychiatric disorders or traits are enriched in differentially accessible peaks during cortical neurogenesis, we calculated partitioned heritability enrichment using LD score regression (Bulik-Sullivan et al., 2015; Finucane et al., 2015a) (Figure 7A). Similar to a previous study in mid-gestation fetal brain (de la Torre-Ubieta et al., 2018), we found cell-type specific enrichments for neuropsychiatric disorders and associated behaviors in regions of open chromatin in fetal brain. Genetic variants associated with several childhood or adult onset neuropsychiatric disorders or traits, including ASD, schizophrenia, major depressive disorder (MDD), neuroticism and depressive symptoms, showed significant partitioned heritability enrichment in peaks more accessible in neural progenitors. With the exception of schizophrenia, these disorders and traits did not show significant enrichment in peaks more accessible in neurons. We observed partitioned heritability enrichment for both intelligence and educational attainment within peaks differentially accessible in both cell types. As a negative control, we did not observe enrichment of inflammatory bowel disease (IBD) heritability in differentially accessible peaks, as expected. These results are consistent with the model (de la Torre-Ubieta et al., 2018; Walker et al., 2019) that genetic variants alter the function of REs during cortical neurogenesis, which then leads to risk for neuropsychiatric disorders or related traits in childhood or adulthood.

**Figure 7:**
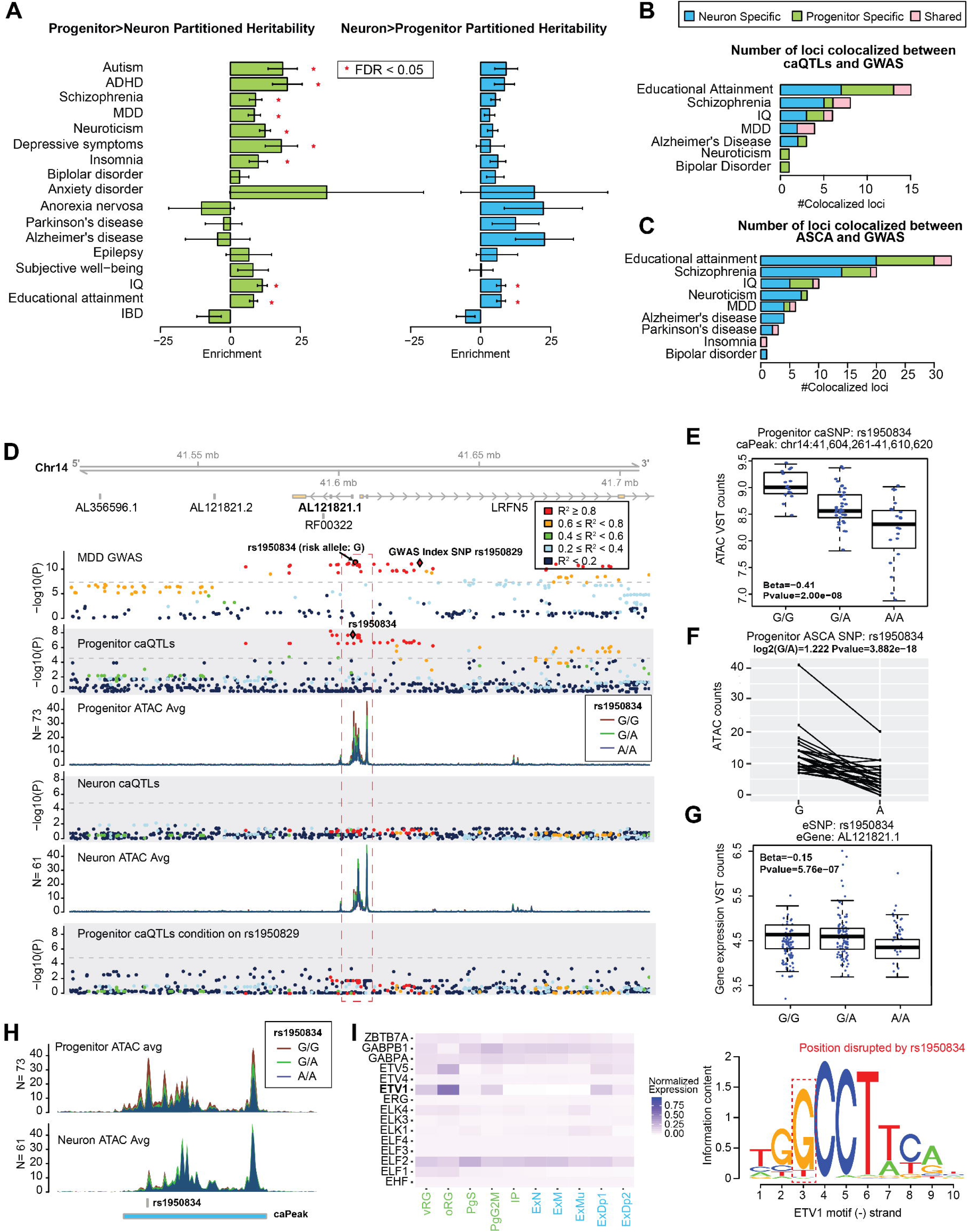
Cell-type specific caQTLs lead to regulatory mechanisms underlying GWAS loci. (A) Partitioned heritability enrichment demonstrated a significant (FDR < 0.05) enrichment of heritability for ASD, Attention Deficit Hyperactivity Disorder (ADHD), schizophrenia, Major Depressive Disorder (MDD), neuroticism, depressive symptoms and insomnia within progenitor > neuron peaks. There was a significant enrichment of heritability for IQ and educational attainment, but not for inflammatory bowel disease (IBD) within progenitor > neuron peaks and neuron > progenitor peaks. (B) Numbers of colocalizations between caQTLs and GWAS loci. (C) Numbers of colocalizations between ASCA and GWAS loci. (D) A colocalized locus between progenitor-specific caQTL and MDD GWAS. (E) Association between rs1950834 and chromatin accessibility of the labeled peak in progenitors. (F) ASCA of rs1950834 in progenitors. (G) Association between rs1950834 and expression of lncRNA AL12182.1. (H) Zoomed in plot of caPeaks colored by genotype at rs1950834. (I) The expression of TFs in which motifs are disrupted by rs1950834 (*left*). The motif logo of ETV1 disrupted by rs1950834 (*right*).

To study the cell-type specific gene-regulatory impact of specific genetic variants associated with these neuropsychiatric disorders or traits, we performed a colocalization analysis of caQTLs in progenitors and neurons with existing GWAS data. We identified putatively co-localized signals (pairwise LD r^2^ > 0.8 between the GWAS index and caQTL index) and then performed conditional analysis to verify that the two variants mark the same locus (Methods). We found co-localized loci in several neuropsychiatric disorders, including schizophrenia (N=8), major depressive disorders (N=4), Alzheimer’s disease (N=3), neuroticism (N=1) and bipolar disorder (N=1), as well as IQ (N=6) and educational attainment (N=15) (Figure 7B). We also found additional ASCA located in GWAS loci (N_Neuronpyschiatric disorders_=83) (Figure 7C). These results indicate that SNPs impact risk for these diseases by regulating the activity of REs in these two cell types during mid-fetal brain development, and provide a framework for exploring the mechanistic bases for these specific disease-associated risk loci.

Next, we investigated regulatory mechanisms underlying co-localized loci using cell-type specific caQTLs. Combining fetal cortical eQTL data, we found a co-localized locus across progenitor specific caQTLs, fetal cortical eQTLs and MDD GWAS (Figure 7D). We found more than 30 SNPs in high LD with an MDD GWAS index SNP (rs1950826). Eight of these variants were located in a caPeak (chr14:41,604,261-41,610,620). We prioritized one putatively causal SNP among those 8 by testing for ASCA, finding that the A allele of the caSNP rs1950834 (protective allele for MDD), was associated with decreased accessibility of this caPeak in progenitors (Figure 7F-7G). We also found rs1950834 was a fetal cortical eSNP, of which the A allele was associated with decreased expression of lncRNA AL121821.1 (ENSG00000258636) in fetal cortex (Figure 7E). After conditioning on the MDD index SNP, rs1950826, the caQTL was no longer significant, indicative of co-localization (Figure 7D). We found evidence to support that this SNP disrupts the binding of ETV1 (Figure 7I; Methods; (Polioudakis et al., 2019)). This suggests that the protective mechanism of this locus for MDD is via the protective allele at the caSNP disrupting binding of ETV1 at a RE in progenitors, decreasing chromatin accessibility of this caPeak, and resulting in decreased expression of lncRNA AL121821.1.

Another example is the co-localized locus between a neuron specific caQTL and schizophrenia GWAS (Supplemental Figure 7A). We found the C allele (schizophrenia protective allele) of the caSNP, rs9930307, was associated with decreased chromatin accessibility of a caPeak (chr16:9,805,171-9,805,430) in neurons (Supplemental Figure 7B). This caSNP was also a neuron-specific ASCA site, providing further evidence of this allele’s impact on chromatin accessibility (Supplemental Figure 7B-7C). After conditioning on the schizophrenia index SNP (rs7191183) in the caQTL analysis, the caSNP was no longer significant (Supplemental Figure 7A). Two possible TF motifs, IRX2 and TP53, were disrupted by this caSNP (Supplemental Figure 7D). Here we prioritize TP53 because TP53 exhibited increased expression in progenitors and maturing neurons. The C allele of rs11643173, a LD proxy of this caSNP (R^2^_rs11643173-C:rs9930307-C_=0.865 in fetal cortical eQTLs), was significantly (FDR < 0.05) associated with increased expression of lncRNA AC022167.1 that is ∼952kb from the caPeak. This data suggests that a possible mechanism leading to risk for schizophrenia is that the risk allele at the caSNP disrupted TP53 binding in a neuron RE, affecting chromatin accessibility of this caPeak, resulting in expression level differences of AC022167.1, with downstream impacts on risk for schizophrenia. We also found a co-localized locus between a progenitor specific caQTL and neuroticism GWAS (Supplemental Figure 7E). The T allele (associated with increased neuroticism) of the caSNP, rs56980069, was significantly associated with increased chromatin accessibility of the caPeak (chr7:38,880,481-38,881,330) in progenitors (Supplemental Figure 7F), but not in neurons. After conditioning on the neuroticism index SNP, rs10244302, the caQTL was no longer significant, indicating a co-localization (Supplemental Figure 7E). We found that the T allele of the caSNP, rs56980069, is associated with disruption of the ZNF713 motif (Supplemental Figure 7G). Using these data, we hypothesize that the neuroticism risk allele disrupted binding of ZNF713, which changes chromatin accessibility of the RE in progenitors.

## Discussion

Our caQTL analysis identified regulatory mechanisms underlying risk variants for neuropsychiatric disorders, cortical structure, and other brain-relevant traits (Supplemental Table S7). The function of genetic variants associated with these traits are often difficult to interpret because most of these variants are mapped to non-coding regions in the genome (Tak and Farnham, 2015). Currently, the function of individual non-coding brain-relevant risk loci is understood through co-localization with eQTLs in adult post-mortem brain tissue or chromatin interaction (Wang et al., 2018; Won et al., 2016). Our study and dataset is able to complement this previous work in several ways: (1) caQTL analysis allows fine mapping of causal variants within correlated LD-blocks by identifying putatively causal variants within peaks; (2) cell-type specific caQTL analysis can prioritize cell-types mediating the risk for neuropsychiatric illness because genetic effects on REs are highly specific; (3) most previous eQTL studies have been performed in post-mortem adult brain cortex (Dobbyn et al., 2018; GTEx Consortium et al., 2017; Wang et al., 2018), but cell-types contributing to the heritability for multiple disorders and traits are not present at this developmental stage suggesting that temporal specificity matters for understanding risk for these disorders (Figure 7; (de la Torre-Ubieta et al., 2018)); and (4) integration of caQTL, eQTL, and brain-trait GWAS allows a more complete understanding of the gene regulatory mechanism leading to risk for neuropsychiatric disorders, where non-coding genetic variants disrupt TF binding to REs, affecting chromatin accessibility, influencing expression of downstream genes, leading to downstream risk for neuropsychiatric disorders. Our study provides a resource to understand the impact of genetic variation on gene regulation during human cortical neurogenesis and provides an additional layer of information to explain the function of common variants associated with risk for neuropsychiatric illness and brain-related traits.

There are several existing methods to map REs in the non-coding genome to the genes they regulate, including chromatin interaction, peak-peak correlation, and high-throughput CRISPRi screens (Corces et al., 2018; Fulco et al., 2019; Won et al., 2016). Here, we integrate caQTL and eQTL studies as an additional method to both identify the genes regulated by REs and to propose causal variants altering the function of these REs. This is complementary to chromatin interaction assays, such as Hi-C, which demonstrate physical interaction with a promoter region, but are not always indicative of regulation (Alexander et al., 2019; Benabdallah et al., 2019).

caQTL and eQTL studies require genetic variation to be present in a population in order to detect their impact. Strong negative selection for genetic variation within REs of genes critical to life may lead to the absence of these variants in the population sampled. Therefore, RE-gene mapping would not be possible for these REs through the integration of caQTL/eQTL datasets. However, for the class of common genetic variation, which is demonstrated to have a large impact on risk for neuropsychiatric disorders and brain traits in aggregate (Sullivan and Geschwind, 2019), this method is useful for both fine mapping and identifying genes regulated by REs.

We provide evidence to support that caQTLs both have higher effect sizes and more cell-type specificity than eQTLs. This suggests that there are a limited number of mechanisms whereby sequence variation impacts chromatin accessibility (caQTL), including TF binding to DNA, whereas there are considerably more mechanisms by which sequence variation can impact transcript levels (eQTL), such as altering TF binding, impacting methylation, or altering miRNA binding sites (Bell et al., 2011; Chen and Rajewsky, 2006; Gaffney et al., 2012). This also suggests that caQTL analyses will identify more genetic variants involved in gene regulation than eQTLs given a limited sample size. Future cell-type specific caQTL analyses will be better powered to identify cell-types involved in the mechanism of GWAS risk variants. However, because our comparison was conducted between cell-type specific caQTLs and tissue-type specific eQTLs, cell-type specific eQTL studies will be necessary to confirm whether the higher effect sizes are caused by stronger genetic impacts on chromatin accessibility or cell-type specificity.

We found that genetic variants associated with differences in chromatin accessibility often disrupted the motifs of TFs involved in neurogenesis, which supports the hypothesis that genetic variation affects chromatin accessibility by disrupting the binding of TFs (Behera et al., 2018). Moreover, we identified which TF motifs were disrupted by genetic variants and found that disruption of motif binding generally leads to decreased chromatin accessibility, with the exception of repressive TFs. These results support a model where chromatin accessibility is increased when activating TFs are bound (Janicki et al., 2004). Our results also support a model where binding of repressive TFs condenses chromatin in REs (Kornberg and Lorch, 1999).

caQTL analysis is able to identify genetic variants associated with REs and to prioritize causal variants, but cannot be used directly to predict the genes regulated by these elements. Most caQTLs did not result in changes in gene expression in bulk fetal cortical tissue. Previous work has suggested that the motif-disrupting alleles do not result in changes to gene expression without the presence of additional transcription factors binding to the same RE (Alasoo et al., 2018). These additional transcription factors may be translocated to the nucleus in response to an external stimulus. caQTLs may therefore be more likely to co-localize with risk loci even in the absence of external stimuli (context-independence), whereas eQTLs would require additional stimuli (context-dependence). This also suggests that future work identifying caQTL/eQTLs in response to environmental stimuli relevant to neural proliferation, differentiation, or function will be especially useful to interpret GWAS risk loci, which is optimally performed in vitro under a controlled setting.

## Acknowledgments

This work was supported by NIH (R00MH102357, U54EB020403, R01MH118349, R01MH120125), Brain Research Foundation, and NC TraCS Pilot funding to JLS. DHG was supported by NIH (5R37 MH060233, 5R01 MH094714, and 1R01 MH110927). The following core facilities were utilized for this project: UNC Neuroscience Center Microscopy Core (P30NS045892), UNC Mammalian Genotyping Core, CGIBD Advanced Analytics Core (NIH grant P30 DK034987), UNC Flow Cytometry Core Facility, UNC Vector Core, UNC Research Computing. Additional core facilities utilized for this project were: UCLA CFAR (5P30 AI028697), and the UCLA Neuroscience Genomics Core. The eQTL data used for the analyses described in this manuscript were obtained from: [https://gtexportal.org/home/] the GTEx Portal on 04/14/19 (v7). The Genotype-Tissue Expression (GTEx) Project was supported by the Common Fund of the Office of the Director of the National Institutes of Health, and by NCI, NHGRI, NHLBI, NIDA, NIMH, and NINDS. We thank Dr. Karen L. Mohlke for helpful comments.

## Author Contributions

JLS, DHG, and LTU conceived the study. JLS provided funding to support the study. ALE, KEC, KPC, MY, LTU, and JLS cultured human neural progenitor cells. ALE performed library preparation. MJL pre-processed the RNA-seq data for eQTL. NA performed eQTL analysis. OK performed immunocytochemistry. MEG, AAK, GEC provided access to adult dlPFC caQTL data. MIL aided in ASCA methodology. DL performed pre-processing, differential accessibility, caQTL, ASCA, co-localization, and motif analyses. JLS and DL wrote the manuscript. All authors commented on and approved the final version of the manuscript.

## Supplemental information

**Supplemental Figure 1: Related to Methods.**
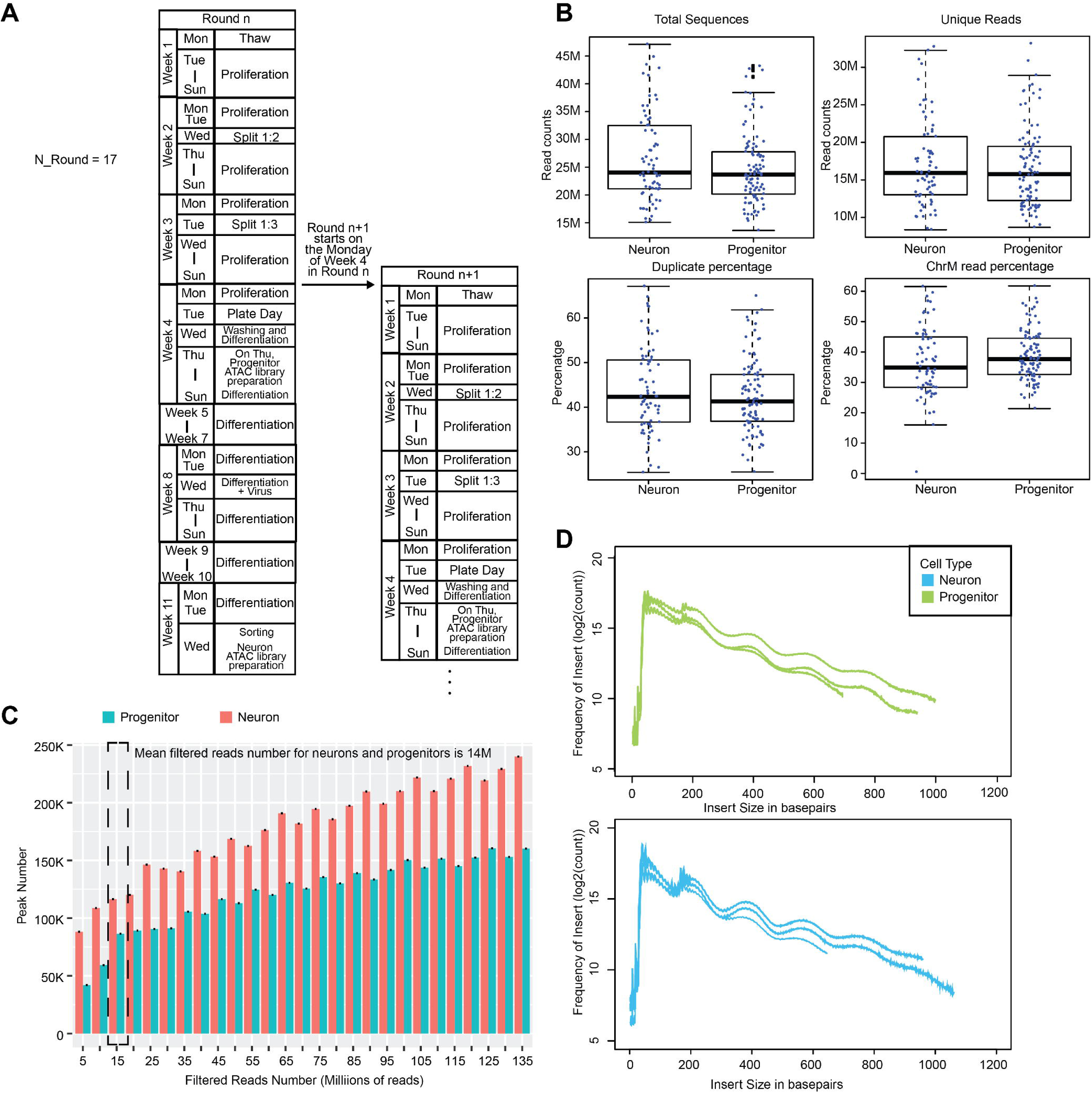
Flowchart for cell culture and pre-processing of ATAC-seq data. (A) Flowchart of cell culture for 17 rounds. (B) Box plot for total sequence depth, unique read number, duplicate percentage, and mitochondria (chrMT) reads percentages in neurons and progenitors. (C) Peak calling versus library sequencing depth. We observed a slower rise in the number of new peaks called after 15 millions filtered read pairs in both neurons and progenitors. This indicates a reasonable balance between read depth and number of new peaks called using an average of 14 filtered million read pairs acquired in our samples. (D) Insert size histograms from 3 randomly selected neuron samples and progenitor samples, showing the expected phasing pattern of transposase insertion between nucleosomes.

**Supplemental Figure 2: Related to Figure 1 and Methods.**
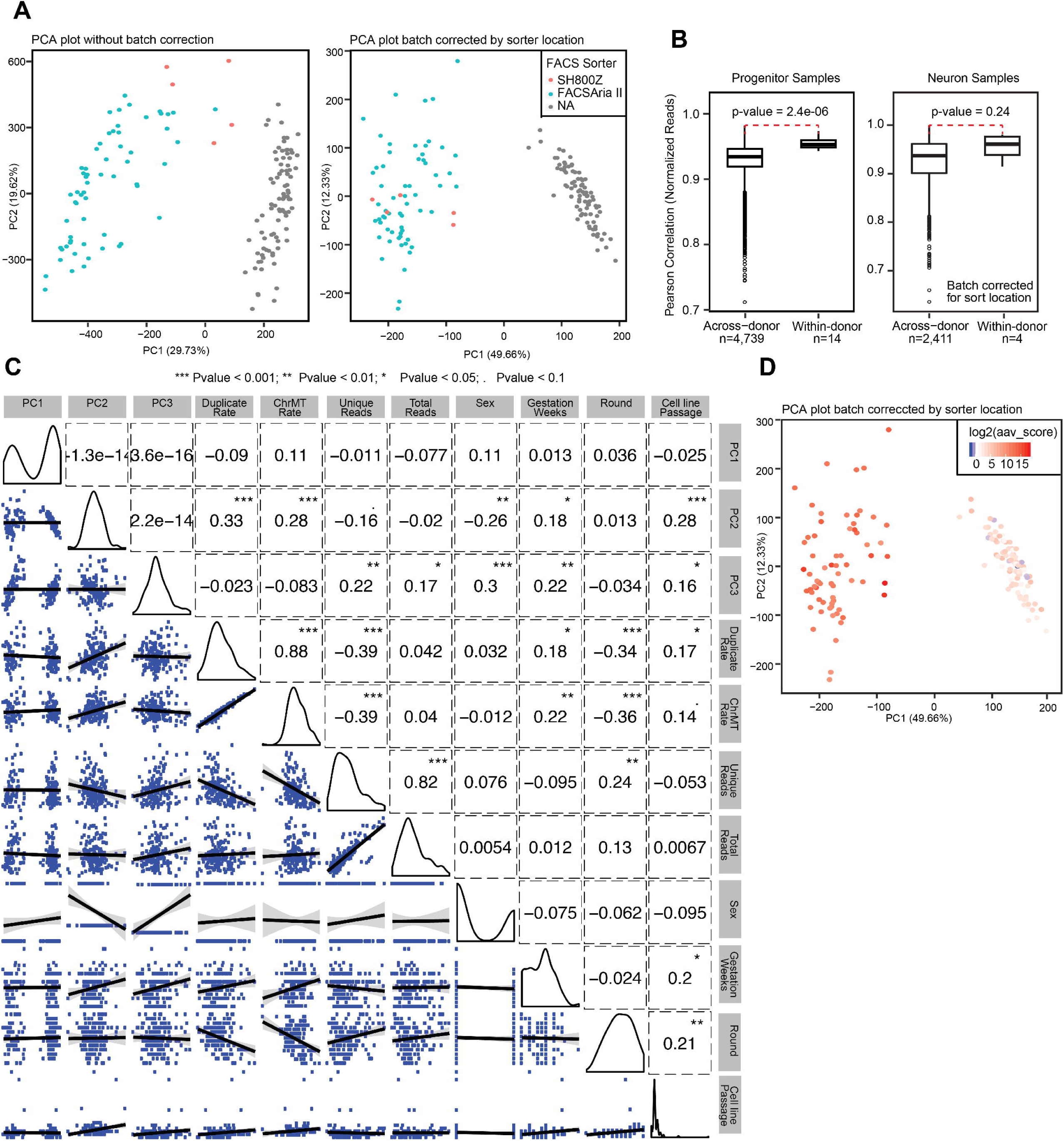
Correlations between PCs from ATAC-seq data and known technical factors. (A) PCA plot for ATAC-seq data before batch correction (*left*) and after batch correction (*right*), colored by sorter. We corrected normalized reads within ATAC-seq peaks in neurons by sorter locations. Then, we corrected normalized reads within ATAC-seq peaks in neurons and progenitors by cell culture round. (B) Correlations of batch corrected normalized reads across donors and within donors in neurons and progenitors. Correlations within donors was significantly higher than correlations across donors in progenitor (p-value=2.4e-06). Correlations within donors was higher than correlations across donors in neurons, but not significant (p-value=0.24), likely due to the fewer number of replicates. (C) Correlations between PC1, PC2 and PC3 from batch corrected normalized reads in neurons and progenitors with known technical and biological factors. (D) PCA plot for batch corrected normalized reads in neurons and progenitors, colored by AAV score. We calculated the AAV score for each sample as the (rate of reads mapped to GFP gene)*1e+08. AAV was not added to progenitor cultures so as expected there are virtually no reads mapping to progenitors. Differences in AAV score among neuron cultures may be indicative of cell health as more infections per cell will occur when there are fewer cells surviving differentiation.

**Supplemental Figure 3: Related to Figure 1 and Methods.**
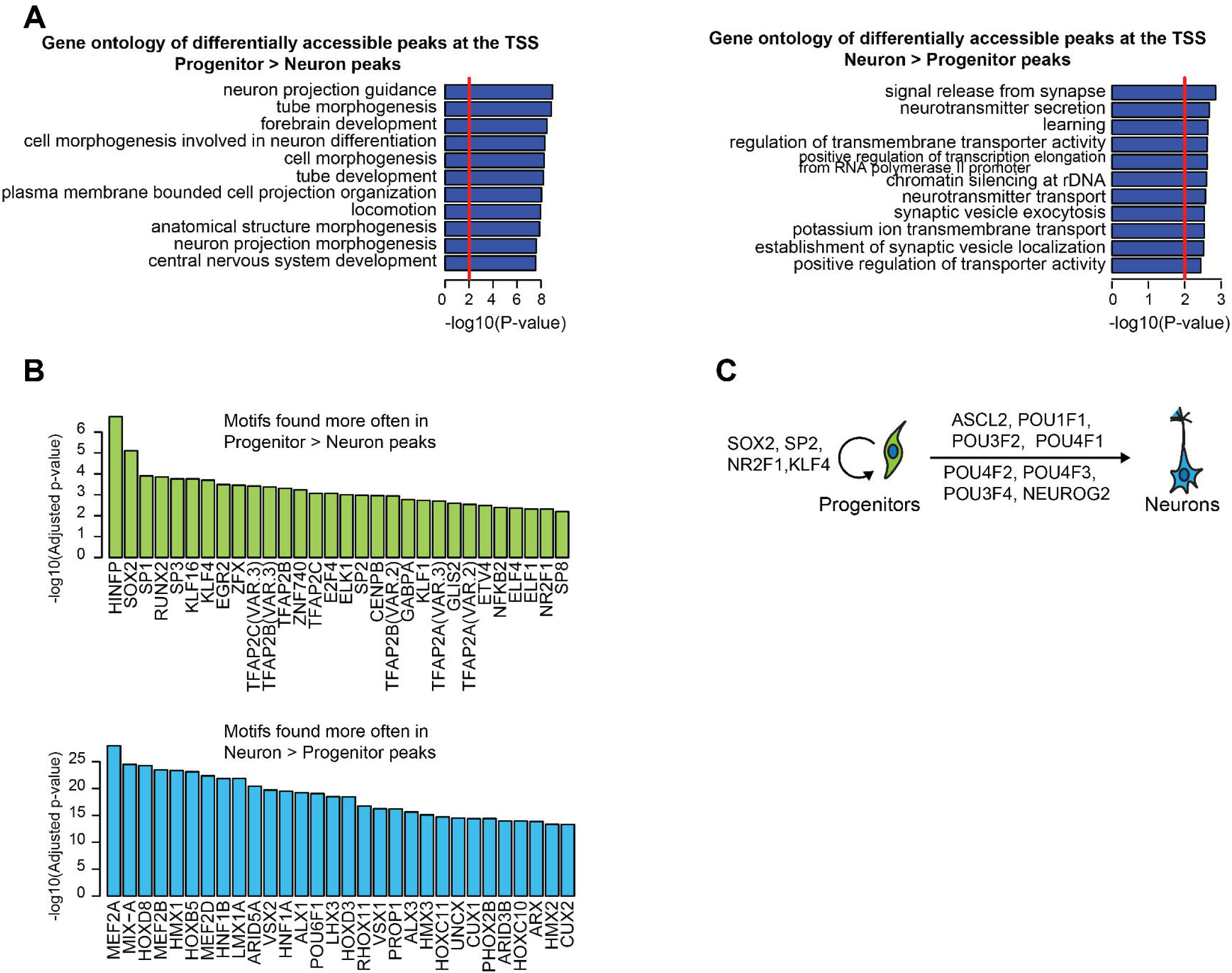
Annotating differentially accessible peaks during neuronal differentiation. (A) Gene ontology (GO) enrichment of differentially accessible peaks at TSS. progenitor > neuron peaks (*left*) and neuron > progenitor peaks (*right*) showed enrichment for GO terms related to proliferation and differentiation, as expected. (B) TFs with significantl differentially enriched conserved binding sites in differentially accessible peaks. The statistical test identifies TFs likely involved in neural progenitor proliferation and maintenance (progenitorTFs; left) or neurogenesis and maturation (neuronTFs; right). The top 30 significantly enriched TFs were shown in this figure, and the full list can be found in Table S2. (C) Schematic of the functions for selected progenitorTFs and neuronTFs.

**Supplemental Figure 4: Related to Figure 2.**
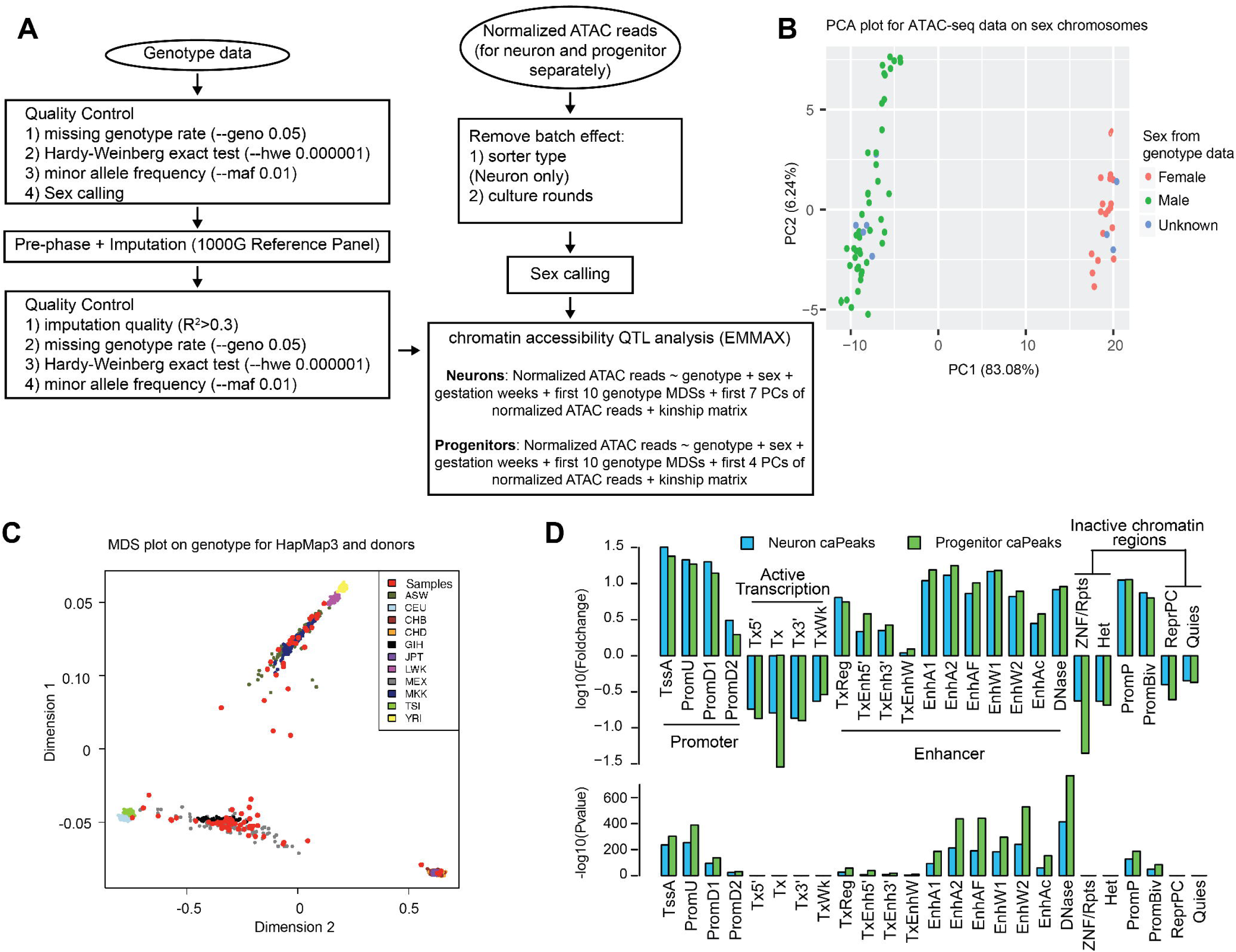
Features of caQTLs. (A) Flowchart for caQTL data analysis. (B) PCA plot for ATAC-seq data on sex chromosomes, colored by sex from genotype data, showing sex could be called using ATAC-seq data. (C) MDS plot for genotype data of HapMap3 and donors in this study, colored by populations from HapMap3 data. ASW: African ancestry in Southwest USA; CEU: Utah residents with Northern and Western European ancestry from the CEPH collection; CHB: Han Chinese in Beijing, China; CHD: Chinese in Metropolitan Denver, Colorado; GIH: Gujarati Indians in Houston, Texas; JPT: Japanese in Tokyo, Japan; LWK: Luhya in Webuye, Kenya; MEX: Mexican ancestry in Los Angeles, California; MKK: Maasai in Kinyawa, Kenya; TSI: Toscans in Italy; YRI: Yoruba in Ibadan, Nigeria. (D) Neuron and progenitor caPeaks enrichment at epigenetically annotated regulatory elements from fetal brain (Epigenetics Roadmap ID = E081).

**Supplemental Figure 5: Related to Figure 4.**
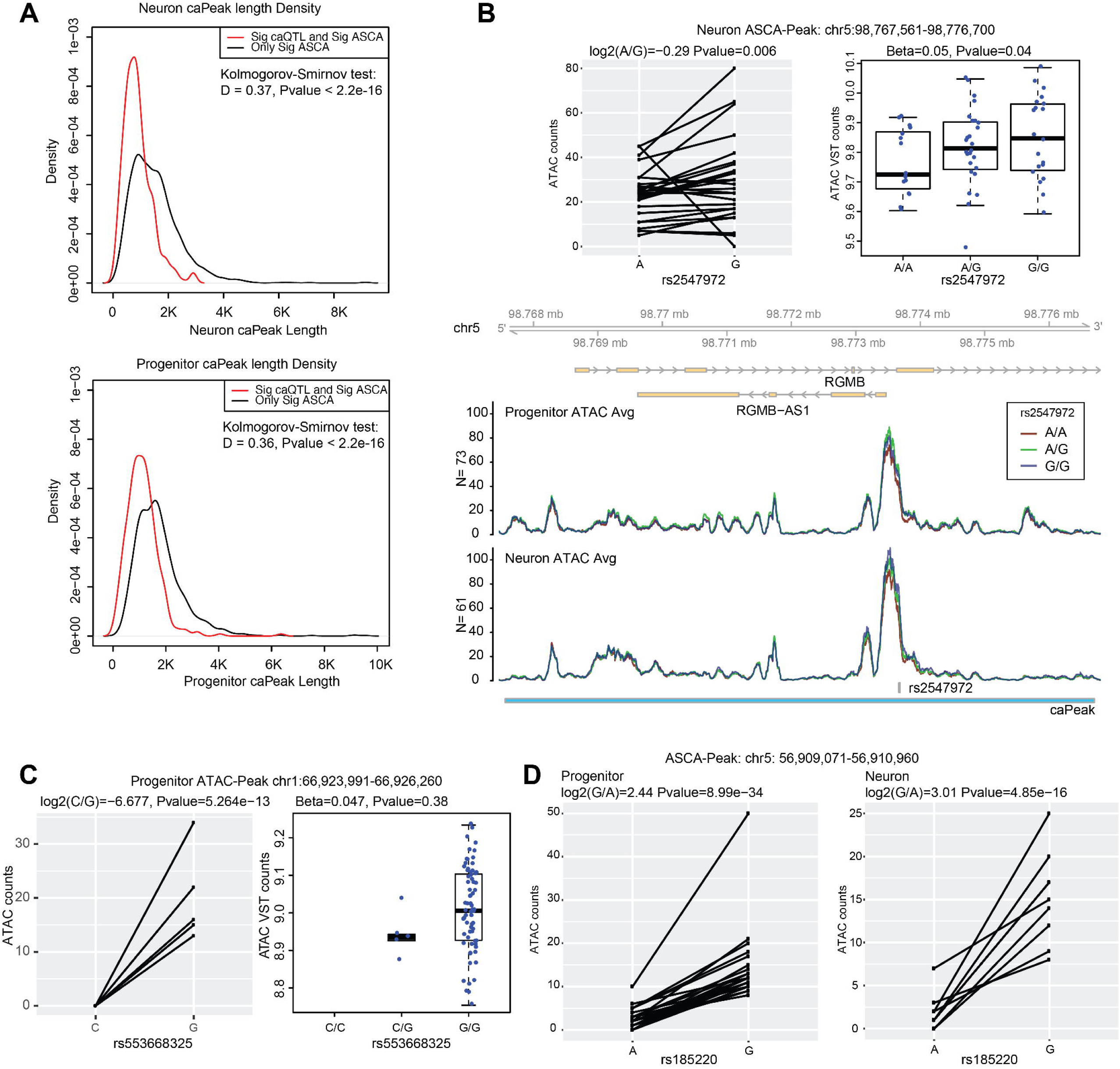
Features of ASCA. (A) Density plot for caPeak length from shared caQTLs and ASCA and from peaks only significant in ASCA in neurons (*top*) and progenitors (*bottom*). caPeak length from only significant in ASCA was significantly larger than caPeak length from shared significant caQTLs and ASCA. (B) The neuron ASCA (caSNP: rs2547972; caPeak: chr5:98,767,561-98,776,700) is not a significant caQTL in neurons because the caPeak was very wide (9,139bp) and only the region near the ASCA SNP shows an association with genotype. (C) The progenitor ASCA (caSNP:rs553668325; caPeak:chr1:66,923,991-66,926,260) is not a significant caQTL in progenitors due to low minor allele frequency leading to less power to detect a caQTL. (D) ASCA between rs185220 (see Figure 3) and chromatin accessibility in progenitors (left) and neurons (right).

**Supplemental Figure 6: Related to Figure 5.**
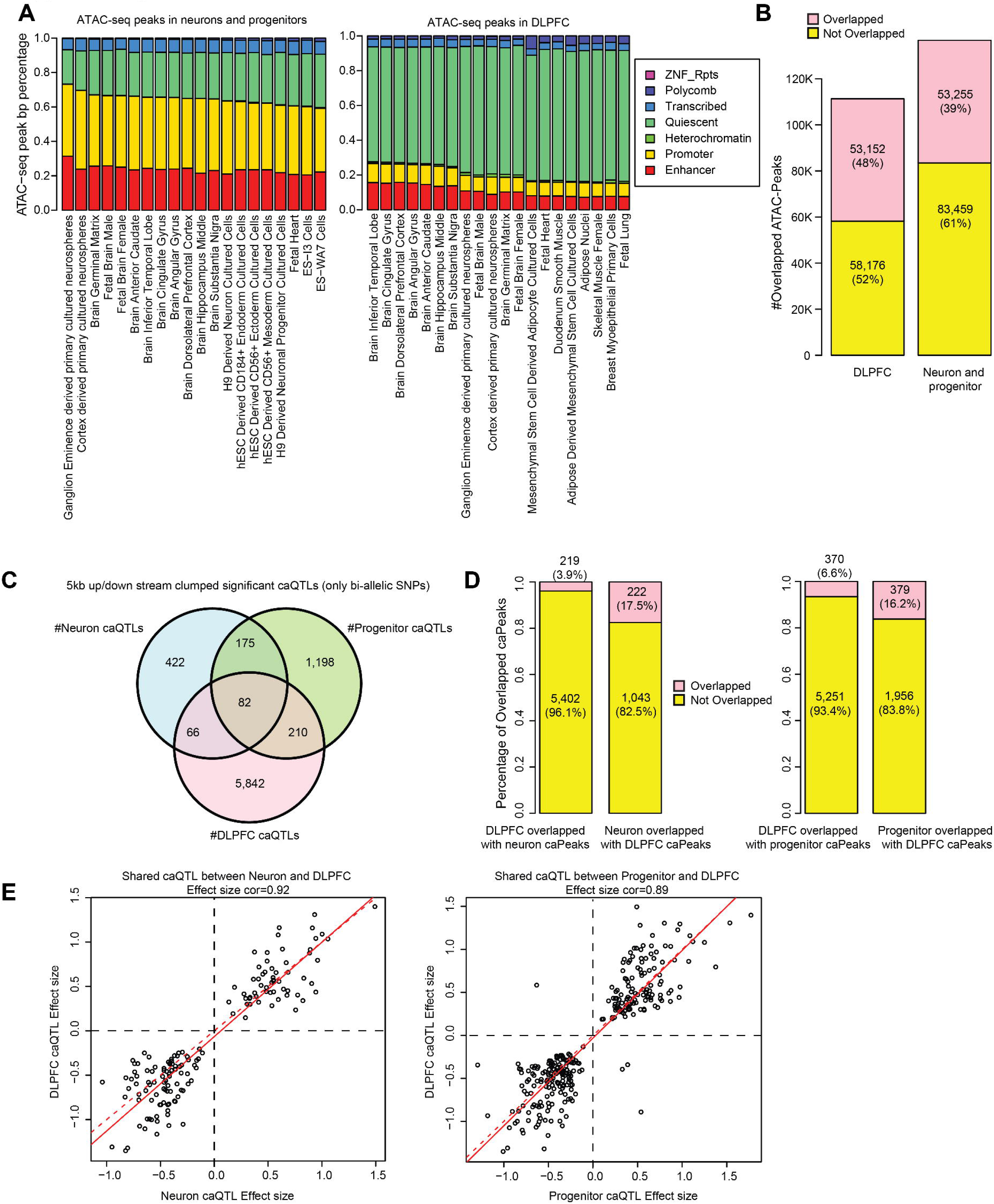
Comparison to adult dorsolateral prefrontal cortex (DLPFC) caQTLs. (A) Accessible peaks (left: neurons and progenitors; right: DLPFC) overlap at epigenetically annotated regulatory elements from different tissues/cell types. Accessible peak bp percentage overlapped with epigenetically annotated regulatory elements. From left to right, tissues ordered by bp percentage overlap with enhancers and promoters. The most enriched tissue for neurons and progenitors accessible peaks are Brain Germinal Matrix and Fetal Brain Female, and the most enriched tissue for DLPFC accessible peaks are Brain Cingulate Gyrus and Brain Dorsolateral Prefrontal Cortex, indicating the accessible peaks are enriched in tissue-specific active regulatory elements. (B) Numbers and percentage of overlapping accessible peaks between cultured cells (neurons and progenitors) and DLPFC. (C) Numbers of overlapped caQTLs between cultured cells (neurons and progenitors) and DLPFC. Here we used clumped caQTLs in 5kb up/down stream from centers of caPeaks (only keeping bi-allelic SNPs). We observed only a very small proportion of overlap, indicating temporal specificity of caQTLs. (D) Numbers and percentage of overlapping caPeaks between neurons/progenitors and DLPFC. caPeaks were temporally and cell-type specific. (E) Correlations of effect sizes for overlapped caQTLs between neurons (*left*) and progenitors (*right*) with DLPFC.

**Supplemental Figure 7: Related to Figure 7.**
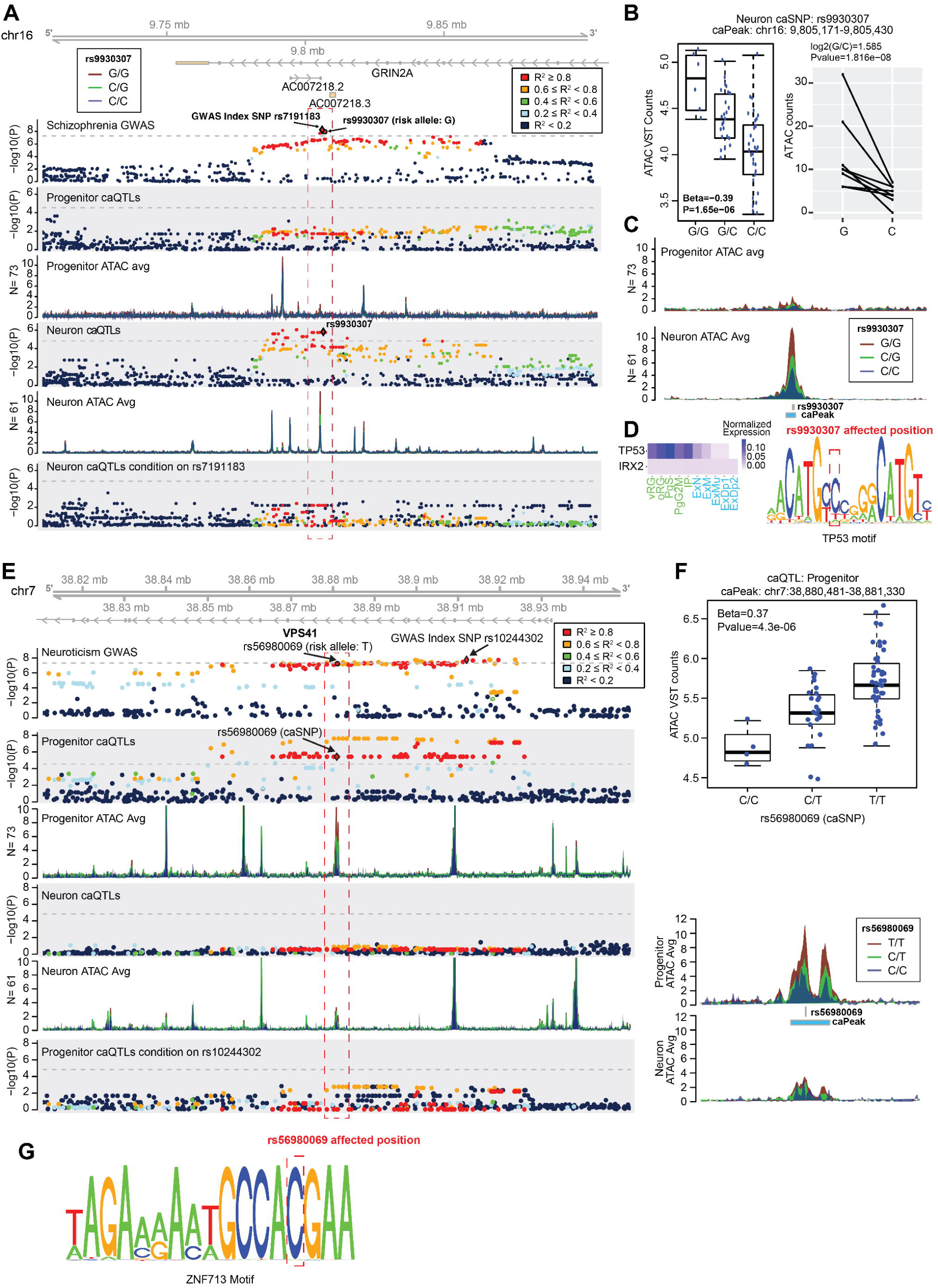
Two examples of cell-type specific caQTLs leading to regulatory mechanisms underlying GWAS loci. (A) The neuron-specific significant caQTL (caSNP: rs9930307; caPeak: chr16: 9,805,171-9,805,430) co-localized with schizophrenia GWAS locus (Index SNP: rs7191183). (B) Box plot for the caQTL (*left*) and ASCA (*right*) (caSNP: rs9930307; caPeak: chr16: 9,805,171-9,805,430). (C) Coverage plot for average ATAC-seq reads for the caPeak (chr16: 9,805,171-9,805,430). (D) Expression of TF motifs, IRX2 and TP53, were disrupted by rs9930307 (*left*). The motif logo of TP53 and the position disrupted by rs9930307. (E) The progenitor-specific significant caQTL (caSNP: rs56980069; caPeak: chr7: 38,880,481-38,881,330) colocalized with neuroticism GWAS locus (Index SNP: rs10244302). (F) Box plot for the caQTL (caSNP: rs56980069; caPeak: chr7: 38,880,481-38,881,330). (G) The motif logo of ZNF713 and the position disrupted by rs1950834.

## Supplementary Tables

**Table S1. Related to Figure 1.** Statistics for differential accessibility at each peak. Seqnames, start, end are the genomic locations of the peaks on hg38; logFC is a measure of effect size between progenitors and neurons with progenitor > neuron corresponding to LFC>0; AveExpr is the value of average reads within peaks; t is the t-statistic, P.Value is the nominal p-value of differential accessibility; adj.P.Val is the Benjamini-Hochberg FDR adjusted p-value; B is B-statistic from limma (Ritchie et al., 2015).

**Table S2 Related to Figures 1.** Significantly enriched TF motifs in differential accessible peaks. TFname and TFID are names of TFs acquired from JASPAR2016. Version is the version for the TF motifs in JASPAR2016. pval, estimate, and Padj are outputs from the differential motif enrichment analysis logistic regression evaluating whether motifs are present more often in progenitor > neuron as compared to neuron > progenitor peaks (estimate > 0; progenitorTF) or present more often in neuron > progenitor as compared to progenitor > neuron peaks (estimate < 0; neuronTF). P-value adjustments for multiple comparisons are via FDR.

**Table S3. Related to Figure 2.** Clumped neuron and progenitor caQTLs. SNP is the caSNP name. Seqnames, peakstart, peakend and PeakID are the genomic locations of the caPeaks on hg38. Distance is the distance (how many base pairs) between the index caSNP and the caPeak center. Beta and Pval are outputs from caQTL analysis, representing effect size and nominal p value. InPeak presents if the caSNP is located in the caPeak. A1 is the non-effect allele and A2 is the effect allele. Rsid is the id for the SNPs.

**Table S4. Related to Figure 2.** Overlapped QTLs between neuron/progenitor caQTLs and fetal cortical eQTLs. SNP is the caSNP name. Chr, peakstart and peakend are the genomic locations of the caPeaks on hg38. caBeta and caP are outputs from caQTL analysis, representing effect size and nominal p value. eGene and hgnc_symbol are names of the eGene. eBeta and eP are outputs from eQTL analysis, representing effect size and nominal p value. eQTL_EffectAllele is the effect allele of eQTL. caQTL_EffectAllele is the effect allele of caQTL. Rsid is the id for index caSNPs.

**Table S5. Related to Figure 4.** Allele Specific Chromatin Accessibility in Neurons and Progenitors. VariantID and SNP are the names of the caSNP. RefAllele and altAllele are reference allele and alternative allele of the caSNP. baseMean is the mean of normalized counts of this SNP for all samples in the analysis; LFC is a measure of effect size between reference allele and alternative allele with altAllele>refAllele corresponding to LFC>0; lfcse is the standard error of LFC for ASCA; stat is the Wald statistic for ASCA; pvalue is the nominal p-value of differential accessibility for ASCA; padj is the Benjamini-Hochberg FDR adjusted p-value for ASCA; C_Het_donors is the number of heterozygous donors, seqnames, start and end are the genomic locations of the caPeaks in hg38. logFC, AveExpr t, P.Value, adj.P.Val and B are from differential accessibility results in Table S1. CaQTL_Beta and caQTL_P are outputs caQTL analysis, representing effect size and nominal p value. caQTL_A1 and caQTL_A2 are non-effect allele and effect allele for the caQTL. Rsid is the rs id for the caSNP.

**Table S6. Related to Figure 6.** TF motifs affected by caSNPs. Rsid is the ID of the caSNP. Strand is the strand of the caSNP and motif. MotifSeq, MotifStart and MotifEnd are the genomic locations of the motif on hg38. REF and ALT are reference allele and alternative allele of the caSNP. snpPos is the genomic locations of the SNPon hg38. motifPos is the position of the caSNP in the motif. geneSymbol and providerId are the name of this motif. dataSource and providerName are the sources of the motif, which come from MotifDb. seqMatch is the match sequence of the motif. pctRef and pctAlt are normalized relative entropy when reference allele and alternative allele in the motif. scoreRef and scoreAlt are relative entropy when reference allele and alternative allele in the motif. Refpvalue and Altpvalue are p values for motif matching when reference allele and alternative allele in the motif. alleleRef and alleleAlt are MAF for the reference allele and alternative allele. Effect is an output from motifBreakR, representing if the motif is strongly affected by the SNP. caQTLCellType represents if the caSNP is progenitor-specific, neuron-specific or shared between neurons and progenitors.

**Table S7. Related to Figure 7.** Co-localized loci for caQTLs and GWAS.rsid is the rs ID od the caSNP. caSNP is the name of the caSNP. CellType represents if the caQTL is progenitor-specific, neuron-specific or shared between neurons and progenitors. GWAS Trait is the trait for tested GWAS. caPeak(hg38) is the genomic locations of the caPeak on hg38. Closest Gene locus is the gene locus which is closest to the caPeak. Effect Allele is the effect allele for the caQTL. Non Effect Allele is the non-effect allele for the caQTL. GWAS Index SNP is the index SNP of GWAS within 100kb upstream and downstream from the center of the caPeak. caQTL InitialBeta is the effect size of the caQTL. caQTL InitialP is the p value of the caQTL. caQTL condBeta is the effect size of the caQTL condition on GWAS index SNP. caQTL condP is the p value of the caQTL condition on GWAS index SNP. AffectedTFs are the TF motifs affected by the caSNP.

## Methods

### Tissue acquisition and cell culture of phNPCs

Human fetal brain tissue was obtained from the UCLA Gene and Cell Therapy Core following IRB regulations. The tissue is often fragmented during acquisition from the surgical procedure. In the lab of Daniel Geschwind, flat, thin pieces of tissue that have the morphology of developing cortex were selected, and in some cases the tissue was sufficiently intact to be certain of cortical identity. Presumed cortical tissue from 14-21 gestation weeks was dissociated into a single cell suspension, cultured as neurospheres, plated for a low number of passages (2.5 ± 1.8 s.d.) on laminin/fibronectin and polyornithine coated plates, and then cryopreserved as human neural progenitors (HNPs) following our previous work (Stein et al., 2014).

Cryopreserved HNPs were shipped to UNC Chapel Hill after signed material transfer agreement by both institutions. All proliferation, differentiation, sorting, library preparation, and analysis were performed at UNC Chapel Hill. In total, HNPs from 92 donors were cultured.

Donors were thawed in “rounds” of approximately 10 donors, so as to create a manageable workload of cell-culture (Supplemental Figure 1A). Donors were randomly assigned into groups and thawed 3 weeks apart. We performed specific experimental events on the same day of the week and had the same interval of time between events for each round. Experimental events included thawing cells, feeding cells, splitting cells, counting and plating cells, washing cells prior to differentiation, coating plates with attachment factors, adding virus, lifting cells for sorting, sorting, and ATAC-seq library preparation. As much as possible, the same researcher performed the same experimental events. We documented if a different researcher performed an experimental event in the database described below. To determine the impact of different rounds, we cultured cells from the same donors in different rounds as technical replicates, for progenitors (N_donor=11, N_replicate=24, in round 1-7,12 and 13) and neurons (N_donor=4, N_replicate=8, in round 2,6,12 and 13).

We thawed cells on a Monday (Supplemental Figure 1A). HNPs were cultured for 8 days using full feeds of proliferation media (1x proliferation media; see stock preparation and media tables below). On day 9, HNPs were split 1:2 and proliferated with half feeds of proliferation media with twice the concentration of growth factors (2x proliferation media) from day 10 to day 14. On day 15, HNPs were split 1:3 and proliferated with half feeds of 2x proliferation media from day 16 to day 21. On day 22, cells were plated for differentiation onto 8 x 6-well plate wells per donor at a concentration of 42,000 cells/cm^2^ (differentiation library preparation wells). Two x 6-well plate wells were also plated for ATAC-seq preparation of progenitor cells (progenitor library preparation wells) in 1x proliferation media. On day 23, all differentiation wells were washed three times with Neurobasal A and then fed with 1x differentiation media (see media tables below). On day 24, the progenitor cells in proliferation media were lifted with trypsin and ATAC-seq libraries were prepared for progenitors. From day 24 through day 84 cells were half fed every Monday, Wednesday and Friday with 2x differentiation media. Virus for labeling neurons (AAV2-hsyn1-eGFP; https://www.addgene.org/50465/; acquired from the UNC Vector Core; (Thiel et al., 1991)) was added at 20,000 MOI for library preparation wells on day 64. On day 84, cells were lifted using Papain (Worthington) with DNase (Worthington) and sent to cell sorter (BD FACS Aria II or Sony SH800S) to sort for live neurons labeled with GFP. Labeled GFP neurons were collected and aliquoted for immediate ATAC-seq library preparation of the neuron cell-type.

All cultures were visually evaluated and ranked with a subjective measure of cell health before ATAC-seq library preparation. Cell health was based on morphology and growth with the highest rank of 2 (mostly healthy cells by brightfield microscopy) and the lowest ranking (many dead cells) of 0.

#### Proliferation media table

| Proliferation base |  | Proliferation 1x |  | Proliferation 2x |  |
| --- | --- | --- | --- | --- | --- |
| Item | Volume | Item | Volume | Item | Volume |
| Neurobasal A (Life Technologies; 10888-022) | 500ml | Proliferation base | 100ml | Proliferation base | 100ml |
| Primocin (Invivogen; ant-pm-2) | 100 µg/ml | EGF/FGF (Life Technologies; PHG0313/PHG0023) | 20 ng/mL | EGF/FGF (Life Technologies; PHG0313/PH | 40ng/mL |
|  |  |  |  | G0023) |  |
| BIT 9500 (Stemcell Technologies; 09500) | 10% | LIF (Life Technologies; PHC9481) | 2 ng/mL | LIF (Life Technologies; PHC9481) | 4ng/mL |
| Glutamax (100x) (Life Technologies; 35050061) | 1% | PDGF (Life Technologies; PHG1034) | 20 ng/mL | PDGF (Life Technologies; PHG1034) | 40ng/mL |
| Heparin (Sigma-Aldrich; H3393-10KU) | 1 µg/mL |  |  |  |  |

#### Differentiation media table

| Differentiation base |  | Differentiation 1x |  | Differentiation 2x |  |
| --- | --- | --- | --- | --- | --- |
| Item | Volume | Item | Volume | Item | Volume |
| Neurobasal A (Life Technologies; 10888-022) | 500ml | Differentiation base | 100ml | Differentiation base | 100ml |
| Primocin (Invivogen; ant-pm-2) | 100 µg/ml | NT-3 (Life Technologies; PHC7036) | 10 ng/mL | NT-3 (Life Technologies; PHC7036) | 20 ng/mL |
| B27 (Life Technologies; 17504-044) | 2% | BDNF (Life Technologies; PHC7074) | 10 ng/mL | BDNF (Life Technologies; PHC7074) | 20 ng/mL |
| Glutamax (100x)<br>(Life Technologies; 35050061) | 1% |  |  |  |  |

#### Plate coating stock table

| Proliferation 1x |  | Differentiation 1x |  |
| --- | --- | --- | --- |
| Item | Concentration | Item | Concentration |
| Fibronectin (Sigma-Aldrich; SLBN9832V) | 5µg/ml | Laminin (Life Technologies; 23017015) | 5µg/ml |
| Poly-L-Ornithine (Sigma-Aldrich; P3655-500MG) | 10µg/ml | Poly-L-Ornithine (Sigma-Aldrich; P3655-500MG) | 10µg/ml |

### Library Preparation for human neural progenitors and neurons

Library preparation was conducted using the published ATAC-seq protocol (Buenrostro et al., 2015). ATAC-seq libraries were prepared immediately following cellular dissociation. Progenitor nuclei were counted using a hemocytometer while neuron nuclei were counted during sorting. 50,000 nuclei were aliquoted into the first step of the ATAC-seq published protocol. Libraries were prepared following the published instructions except that the last clean up step was modified to use KAPA pure beads (AmpureXP beads at a 1:1 ratio to remove dNTPs, salts, primers or primer dimers) instead of Qiagen Minelute clean-up kit. All libraries were sequenced to a minimum depth of 13.6M and an average depth of 25.5M using 50 bp PE sequencing on an Illumina HiSeq2500 machine (Supplemental Figure 1B). In total, we acquired 98 ATAC-seq libraries from progenitors (N_donor_=86, N_libraries replicated_=12) and 70 ATAC-seq libraries from neurons (N_donor_=67, N_libarary replicated_=4). All libraries were sequenced to an average depth of 25.5 (± 7.21 s.d.) million read pairs (Supplemental Figure 1B), which resulted in an average depth of 14 (± 4.8 s.d.) million reads pairs per sample after filtering for mitochondrial contamination and duplicates. We performed a sensitivity analysis for read depth vs peak calling that showed greater than 15 million filtered read pairs per library led to a fewer number of new peaks called, indicating a reasonable balance between read depth and peaks called on the libraries generated here (Supplemental Figure 1C).

### Recording technical variables and randomization

To reduce the impact of batch effects on interpretation of our results, we attempted to either have no batches when possible (e.g., perform all experiments using the same lot of a reagent) or when this was not possible, randomize technical variables (round a donor was thawed in, sequencing pool) such that they had minimal correlation with variables of interest. In order to extensively document the impact of technical variables on outcome measures, we maintained a relational MySQL database which allowed us to keep track of many technical and biological variables throughout each experimental event. Each downstream ATAC-seq library preparation therefore is able to be tracked back to all technical and biological variables associated with its cell culture. The variables recorded were:

*Media*: Basal media lots, growth factor lots, supplement lots, antibiotic lots; *Virus:* Lot number; *Donor:* sex inferred from genotype; gestation week; *Culture:* passage, round, thaw date, each split date, split ratio, trypsin lot, PBS lot, polyornithine lot, fibronectin lot, plate, and well position, cells per well, date of virus addition, differentiation time, date of differentiation media addition, person plating for differentiation, virus used, person performing splits, person performing virus addition, virus lot, virus multiplicity of infection (MOI), laminin lot, dissociation lot, person performing dissociation of neuronal cultures; *Cell sorting:* Sort date and time, number of live cells, number of GFP+ cells, total number of cells, FACS machine; *ATAC-seq library preparation*: number of cells input to the library preparation, person performing the cell lifting, lysis date, PBS lot, lysis buffer batch, Illumina Kit lot, PCR master mix lot, PCR clean up kit lot, number of time pipetting up and down during lysis, number of times pipetting up and down during transposase reaction, transposase reaction volume, barcode indices used for multiplexing of each sample, number of PCR cycles added in the ATAC-seq protocol, final DNA concentration after library preparation complete; *Sequencing:* sequencing date, sequencing company, type of sequencer, read length.

Randomization was performed multiple times. First, randomization was performed to assign each donor to a thawing “round”. Randomization was performed at this stage by randomly ordering all donors and selecting those to go in each round (generally about 10 donors per round). After culture and library preparation were complete, randomization was performed to assign each library preparation to a pool for sequencing. Randomization was performed using custom R code to minimize the correlation of sequencing pool with concentration of the library, barcode index (assuring that no barcodes were represented in more than one pool), cell type (either neuron or progenitor), round cells were cultured in, and donor.

### ATAC-seq data pre-processing

Sequencing reads were first quality controlled via fastqc (v0.11.7) to check for sequence quality in each library. We observed high quality sequencing for all libraries (PHRED > 20, average duplication rate = 43.07% which is almost entirely mtDNA contamination (Supplemental Figure 1B) which is in agreement with previous studies using the same ATAC-seq method (de la Torre-Ubieta et al., 2018), and average GC content = 45%) (http://www.bioinformatics.babraham.ac.uk/projects/fastqc/). Sequencing adapters were removed using BBMAP/BBDUK (https://jgi.doe.gov/data-and-tools/bbtools/bb-tools-user-guide/bbduk-guide/).

Then, sequencing reads were mapped to the human genome including decoy sequences (GRCh38/hg38) using bwa mem (Li, 2013) (v0.7.17). Optical and PCR duplicates were then removed using Picard tools (http://broadinstitute.github.io/picard/faq.html) (v2.18.22). Only uniquely mapped reads mapping to chr 1-22 and X were kept (mitochondrial genome, Y chromosome, and unmapped contigs were removed) using samtools (Li et al., 2009) (v1.9). Sequencing reads mapped to ENCODE blacklist regions (http://hgdownload.cse.ucsc.edu/goldenPath/hg19/encodeDCC/wgEncodeMapability/wgEncodeDacMapabilityConsensusExcludable.bed.gz, converted to hg38 using UCSCtools/liftOver (v320)), were then removed by bedtools (Quinlan and Hall, 2010) (v2.26).

We did a sensitivity analysis for peak calling using pre-processed bam files. It showed acquiring 9x higher read depth resulted in only 70,000 additional peaks by MACS2 (Feng et al., 2011)https://github.com/taoliu/MACS (Supplemental Figure 1C). So we reasoned that our acquired sequence depth obtained a reasonable balance between additional read depth and number of peaks called. We calculated the insert size of pre-processed bam files using Picard tools (v2.18.22). The insert size histogram shows clear periodicity representative of preferential Tn5 binding around nucleosomes (Supplemental Figure 1D).

Peaks were called for all samples using CSAW (Lun and Smyth, 2016) (v1.16.1), which identifies peaks with smaller peak length than previous methods (MACS2 and DiffBind) and showed higher enrichment at active regulatory elements such as enhancer and promoters (data not shown). For CSAW, peaks were identified in 10 bp bins with average read number greater than 5 across all samples (both neurons and progenitors). Bins directly next to each other (1 bp minimum distance) were merged to call a peak. For all samples (N_Neuron=67, N_Progenitor=86), CSAW identified 136,714 peaks.

R v3.4.1 was used for all subsequent analyses. The number of reads within each CSAW-called peak were counted and then normalization factors for each peak across samples were calculated accounting for GC content, peak width, and total number of unique non-mitochondrial fragments sequenced using conditional quantile normalization (Hansen et al., 2012) from the cqn package (v1.28.1). Variance stabilizing transformed (VST) counts were calculated using DESeq2 (Love et al., 2014) (v1.22.2) for batch effect correction and differential accessibility analysis by limma (Ritchie et al., 2015) (v3.38.3).

As two different sorters were used to sort GFP+ neurons (63 neuron cell lines (N_donor_=61) for one sorter and 7 cell lines (N_donor_=5) for another sorters), and we detected that sort location had a strong effect on PCA from neuron samples, sorter locations in neuron VST counts were first corrected (limma batcheffectremove()) (Supplemental Figure 2A). Corrected neuron VST counts and progenitor VST counts were combined so that the potential batch effect from cell culture rounds were corrected (limma removeBatchEffect).

### Mycoplasma contamination checks

To check if there was any contamination from mycoplasma while in culture, we downloaded 98 mycoplasma genomes (from NCBI) and then mapped all ATAC-seq data to every mycoplasma genome. Less than 0.01% of each ATAC-seq sample mapped to any mycoplasma genome, which demonstrated that our cultures were not contaminated with mycoplasma.

### Replicate correlations, principal component analysis, and correlation with technical factors

To determine the reliability of our experiment, we cultured the same donor multiple times. We then correlated the batch corrected VST counts in CSAW peaks for neuron and progenitor replicates either within donors or calculated correlations across donors (Supplemental Figure 2B). The correlations of replicates within donors are higher than that of samples across donors in both neuron and progenitor samples.

The principal component analysis for batch corrected VST counts for all samples was done using the prcomp() R function. Then the correlations for the first 3 PCs and all technical variables that we recorded were calculated using R. The technical variables include sex, gestation week, cell line thaw passage, duplication rates, total sequence number, percentage of reads in chrMT, number of unique reads, sequencing barcode index number, pool for both sequencing and library preparation, healthy rank of differentiated cells, round of cell culture, proliferation media batch, health rank of progenitor cells, PBS lot number, fibronectin lot number, Illumina Kit (Tn5 transposase) lot number, number of unique non-chrMT sequences, papain lot number, individual who added papain, individual who added virus, sequencing lane, sorter locations, individual who plated cells, individual who differentiated cells from progenitors to neurons, the number of differentiated cells prior to sorting, SYN1 promoter coverage (inferred from the number of unique reads mapping to the viral plasmid), and number of reads mapped to GFP gene. We found significant correlation of PC2 with duplication rates (r= 0.33, p<0.001), percentage of reads in chrMT (r=0.28, p < 0.001), sex (r=-0.52, p< 0.001), gestation week (r=-0.23, p< 0.01), cell line thaw passage (r=0.28, p<0.001); PC3 with number of unique reads (r=0.22, p< 0.05), total sequence number (r=0.17, p< 0.05), gestation week (r=0.22, p< 0.01), sex (r=0.3, p< 0.001), and cell line thaw passage (r=0.16, p<0.001) (Supplemental Figure 2C). We calculated the aav_score for each sample using the (rate of reads mapped to GFP gene)*1e+08, and we found neuron samples have much higher aav_score than progenitor samples (Supplemental Figure 2D).

### ATAC-seq differential accessibility analysis

To ensure independence, we randomly selected one library from each donor for each cell-type (technical replicates where one donor was cultured multiple times for a given cell-type) were excluded. In order to find differentially accessible peaks across cell type controlling for technical factors, the dependent variable was the batch corrected number of reads within CSAW peaks and the linear regression model independent variables included a regressor for cell type (neuron or progenitor) and a factor regressor for donor IDs included in the analysis.

### Enrichment of peaks within annotated regions of the genome

Enrichment of differentially accessible peaks within annotated genetic regions of the genome or epigenetically annotated regions of the genome was calculated using the ratio between the (#bases in state AND overlap feature)/(#bases in genome) and the [(#bases overlap feature)/(#bases in genome) X (#bases in state)/(#bases in genome)] as described previously by the Roadmap Epigenomics Consortium (Roadmap Epigenomics Consortium et al., 2015). The significance of this enrichment was calculated using a binomial test as in the GREAT algorithm (McLean et al., 2010).

Chromatin state definitions from an imputed 25-state model were derived from fetal brain tissue (E081) and other *in vivo* tissues/cell types by the Roadmap Epigenomics project (Ernst and Kellis, 2015; Roadmap Epigenomics Consortium et al., 2015) and acquired from (http://www.broadinstitute.org/~jernst/MODEL_IMPUTED12MARKS/) after liftOver to hg38 (0.001% of peaks could not be lifted over). We generated the following combined states by merging states of similar genomic context: Promoter(2_PromU, 3_PromD1, 4_PromD2, 22_PromP, 23_PromBiv), Enhancer (13_EnhA1, 14_EnhA2, 15_EnhAF, 16_EnhW1, 17_EnhW2, 18_EnhAc), Heterochromatin (21_Het), Quiescent (25_Quies), Transcribed (1_TssA, 5_Tx5’, 6_Tx, 7_Tx3’, 8_TxWk, 9_TxReg, 10_TxEnh5’, 11_TxEnh3’, 12_TxEnhW), Polycomb (24_ReprPC) and ZNF_Rpts (20_ZNF/Rpts).

Locations of ATAC-seq peaks in fetal brain were acquired from previously published work (de la Torre-Ubieta et al., 2018). After liftOver to hg38, 0.001% of peaks could not be lifted over.

Gene ontology analysis at the TSS for differentially accessible peaks (Supplementary Figure 3) was completed by first overlapping differentially accessible peaks with a region 2kb upstream and 1kb downstream of the TSS of genes defined by homo sapiens gene ensembl version 78 GRCh38.p12. Protein-coding genes with promoter overlapped with selected differentially accessible peaks (|LFC|>0.5) were input into the TopGO (Alexa and Rahnenfuhrer, 2010) package (v2.34.0), with all protein-coding genes as background.

Gene based annotations of the genome were derived from Homo sapiens gene ensembl version 78 (GRCh38.p12) for plotting loci.

### Differential transcription factor binding analysis

We performed an analysis to identify motifs with differential prevalence in differentially accessible peaks (Table S2, Figure S3B). To avoid the bias caused by different number of progenitor > neuron and neuron > progenitor differentially accessible peaks, here we only used top 2000 progenitor > neuron peaks with highest LFC, and top 2000 neuron > progenitor peaks with lowest LFC.

Potential transcription factor binding sites were called in the human genome using TFBSTools (v1.4.0) with a minimum score threshold of 80% based on position weight matrices from the JASPAR2016 (Mathelier et al., 2016) core database, selecting vertebrates as the taxonomic group. Only the most recent version of the PWM for a given TF was used. To select regions of the genome that are highly conserved among vertebrates, and likely functional, 100-way phastCons (Pollard et al., 2010) scores > 0.4 in regions ≥ 20 bp were saved (downloaded from UCSC genome browser). Only called TFBS sites within conserved regions were retained for further analyses. Differential motif enrichment analysis was performed using a logistic regression model to identify motifs present more often in progenitor > neuron peaks as compared to neuron > progenitor peaks, or vice versa. Logistic regression explicitly controlled for differences in peak width and peak conservation between progenitor > neuron and neuron > progenitor differentially accessible peaks. The analysis was implemented in R as: glm(TFBS ∼ ProgenitorNeuron + peakwidth + conservedbppercent, family = “binomial”). The dependent variable (TFBS) was a binary representation of whether each differentially accessible peak contained a motif of a TF or not. The independent variable of interest marked whether a peak was progenitor > neuron (ProgenitorNeuron=1) or neuron > progenitor (ProgenitorNeuron=0). Other covariates included peak width (peakwidth) and the percentage of the peak with conservation (conservedbppercent) as defined above. Significant differential motif enrichment was determined by FDR adjusted P-value < 0.05 threshold of the ProgenitorNeuron covariate. progenitorTFs were defined as significant differentially enriched motifs present more often in progenitor > neuron as compared to neuron > progenitor peaks, whereas neuronTFs were defined as significant differentially enriched motifs present more often in neuron > progenitor as compared to progenitor > neuron peaks.

### Genotype pre-processing

Genotyping was performed using Illumina HumanOmni2.5 or HumanOmni2.5Exome platform. SNP genotypes were exported into PLINK format. SNP marker names were converted from Illumina KGP IDs to rsIDs using the conversion file provided by Illumina. Quality control was performed in PLINK v1.90b3 (Chang et al., 2015) (Supplemental Figure 4A). SNPs were filtered based on Hardy-Weinberg equilibrium (--hwe 1e6), minor allele frequency (--maf 0.01), individual missing genotype rate (--mind 0.10), variant missing genotype rate (--geno 0.05) resulting in 1,760,704 directly genotyped variants. Multidimensional scaling (MDS) analysis of genotypes from all individuals used in the study was completed in PLINK v1.90b3. We did not see a strong effect of genotyping batch on genotype data based on MDS1 and MDS2 from different genotyping batches. We used PLINK v1.90b3 to call sex from genotype data. For the sampels with unkown sex from genotype data, we ploted PCA results (PC1 vs PC2) of ATAC-seq reads on sex chromosomes (chromsome X and Y) to identify sex (Supplemental Figure 4B).

### Sample Swap and contamination Identification

Quality controlled genotype data and BAM files were used to identify any sample swaps between the ATAC-seq and genotyping data using VerifyBamID v1.1.3 (Jun et al., 2012). We removed any BAM file with [FREEMIX] > 0.02 (N_donor=4), and corrected sample swaps (N_donor=7). After this filtering step, our sample size was comprised of 73 unique donors for progenitor samples and 61 unique donors for neuron samples for the caQTL studies.

### Imputation

Filtered genotype data were pre-phased by SHAPEIT (Delaneau et al., 2011) v2.837. Minimac4 (Das et al., 2016) (v1.0.0) was used to impute the filtered genotyped markers using reference haplotype panels from the 1000 Genomes Project (The 1000 Genomes Project Consortium Phase 3) that contain a total of 37.9 million SNPs in 2,504 individuals from any ancestry, including those from West Africa, East Asia and Europe. For the variants on chrX, we separated chrX into pseudoautosomal regions and non-pseudoautosomal regions, then pre-phased and imputed them separately.

After genotype imputation, we extracted the genotypes for all individuals assayed for chromatin accessibility. Imputed genotype data were filtered for variant missing genotype rate < 0.05, Hardy-Weinberg equilibrium p-value < 1 x 10^-6^ and minor allele frequency (MAF) 1%. Imputation quality was assessed filtering variants where allele dosage Rsquared > 0.3 by Minimac4, resulting in ∼13.6 million SNPs.

### caQTL mapping

We calculated multidimensional scaling (MDS) for genotype data of our samples and genotype data from HapMap3 (https://www.sanger.ac.uk/resources/downloads/human/hapmap3.html) following the protocol from ENIGMA consortium (http://enigma.ini.usc.edu/wp-content/uploads/2012/07/ENIGMA2_1KGP_cookbook_v3.pdf). We identified multiple ancestries of donors of our samples in the MDS plot (MDS1 vs. MDS2) (Supplemental Figure 4C).

To control for population stratification and cryptic relatedness of our samples when mapping caQTLs, we ran caQTL analysis with EMMAX (Kang et al., 2010), which accounts for population structure using a genetic relatedness or kinship matrix. We used the emmax-kin function (-v –h -s -d 10) to create the IBS kinship matrix for each tested genetic variant from non-imputed genotype data excluding all genetic variants on the same chromosome with the tested genetic variant (Price et al., 2010).

We performed proximal caQTL mapping using a window of 100 kb up- and down-stream of the center of 136,714 csaw peaks using batch corrected read counts of each peak for each donor (Supplemental Figure 4A). We performed caQTL analysis separately in neurons and progenitors using imputed genotype data. To prevent results driven by only one minor allele homozygous donor, we retained the variants with at least 2 minor allele homozygous donors or at least 2 heterozygous donors. In addition to the kinship matrix (Price et al., 2010) we used the following covariates in the association model: gestation week, sex, 10 genotype MDSs. In addition, for the progenitor caQTLs we corrected for 4 PCs across VST counts of the chromatin accessibility data. For neurons, we corrected for 7 PCs of VST counts of the chromatin accessibility data. These numbers were chosen to maximize the number of caQTLs for each cell-type. Nominal EMMAX p-values were corrected for multiple testing using the Benjamini–Hochberg FDR correction (Benjamini and Hochberg, 1995) within neuron caQTLs and within progenitor caQTLs separately (FDR < 0.05). The percent variance explained was calculated using the method from a previous study (Shim et al., 2015).

### Identify correlated caPeaks

To identify correlated caPeaks, we defined primary caPeaks as the caPeaks harboring caSNP(s). We then defined secondary caPeaks as peaks which are associated with the caSNP of a primary peak. We calculated Perason’s correlation between the primary caPeak and all caPeaks within +/− 2Mbp from the center of its secondary caPeak (including the secondary caPeak), then corrected the Pearson’s correlation p-value using the Benjamini–Hochberg FDR correction (Benjamini and Hochberg, 1995). If the secondary caPeak was significantly (FDR < 0.05) correlated with the primary caPeak, this caSNP-caPeak pair was classified as “caSNP in correlated caPeak”.

### Allele specific chromatin accessibility

In order to decrease the impact of mapping bias, we used WASP (2018-07) (van de Geijn et al., 2015) to remap the sequencing reads which intersect with any bi-allelic SNP to the human genome (GRCh38/hg38) using genotype data for each sample and removing the duplicate reads. We then removed reads mapping to the mitochondrial genome, Y chromosome using samtools (Li et al., 2009) (v1.9). Sequencing reads mapped to ENCODE blacklist regions were removed via bedtools (Quinlan and Hall, 2010) (v2.26). We used GATK tools to extract allele specific read counts for every SNP. We first filtered for SNPs within each donor that had sufficient read depth by retaining SNPs with total counts greater than or equal to 10 for neuron and progenitor samples, separately. Then to calculate allele specific chromatin accessibility, we retained those SNPs with average read counts for all heterozygous donors greater than or equal to 15. Finally, we retained only those SNPs that meet these previous thresholds for at least 5 heterozygous donors. DESeq2 was used to calculate the LFC (Alternative read counts/Reference read counts) for filtered SNPs across all heterozygous donors. The non-heterozygous donors were excluded from the differential analysis for each SNP using sample-specific weights, and maximum likelihood estimation was used for dispersion estimation followed by Wald tests of the estimated LFC. FDR < 0.05 was used as the threshold for significance.

### Bulk Fetal brain eQTL mapping

Bulk fetal cortical wall eQTL data described in a previous publication (Walker et al., 2019), was re-analyzed in this study with the following modifications: (1) here we used a linear mixed model implemented in EMMAX to more stringently control for population stratification, and (2) here we add 7 more donors to the analysis because these donors were genotyped after the publication of the previous manuscript. rRNA-depleted RNA-seq data from flash frozen human fetal brain cortical wall tissues derived from 235 donors at 14-21 gestation weeks were used for eQTL analysis. Gene based annotations of the genome were derived from Homo sapiens gene ensembl version 92 (GRCh38) for eQTLs. Only genes which are expressed in more than 5% of donors with at least 10 counts were included in the analysis. VST normalized expression values were used as phenotypes for eQTL analysis. Genomic DNA from human fetal brain cortical wall tissues derived from 235 donors at 14-21 PCW was extracted. Each donor tissue was genotyped on a dense array (Illumina Omni 2.5+Exome) and imputed to a common reference panel (1000 Genomes; described above). Variants were retained in the analysis if there were at least 2 heterozygous donors and no homozygous minor allele donors, or if there were at least 2 minor allele homozygous donors. For the effect size comparison analysis fetal brain eQTL vs caQTLs (Figure 2G), we subsampled fetal brain eQTL donors to the same sample size as the caQTL while maintaining the population composition similar to the larger donor list.

Cis-eQTL analysis was performed by evaluating association between each gene’s expression and variants within ±1 Mb window of transcription start site of each gene by implementing linear mixed model association software, EMMAX (Kang et al., 2008). All markers on the chromosome of this candidate marker were excluded from the IBS kinship matrix was generated with emmax-kin function (-v -h -s -d 10), and added as a random variable into linear mixed model for association test. In addition to kinship matrix, 10 MDS components of genotype, sex, and first 10 PCs of gene expression were included into the covariate matrix. After association, nominal p-values were corrected for multiple testing using the Benjamini Hochberg FDR correction, and associations with lower than 5% FDR threshold value were accepted as significant. In order to obtain LD-independent eQTLs, a clumping procedure was implemented with PLINK v1.90b3 with the parameter --clump-kb 500 --clump-p1 0.0002957020645 (FDR threshold p-value) --clump-p2 0.01 --clump-r2 0.50.

### Identifying LD-independent caQTLs and overlaps between caQTLs and eQTLs

To identify LD-independent caQTLs, we used PLINK v1.90b3 with parameters (--clump-kb 250 --clump-r2 0.50 --clump-p2 0.01), for --clump-r1 we used p values with FDR =0.05 for neuron donor genotypes (p <= 1.49e-05) and progenitor donor genotypes (p <= 2.88e-05) separately. For multiple index SNPs in perfect LD (r^2^=1), we only kept the SNPs that are closest to the ATAC-seq peak center.

In order to determine if caQTLs overlap between progenitors and neurons, we accounted for LD. For caQTLs in progenitor samples and neuron samples, we listed the SNPs with pairwise LD r^2^ > 0.5 with index caSNPs for each shared caPeak using LD matrices from progenitor donors genotype data and neuron donors genotype data, separately. Then for each shared caPeak, if any listed SNP was found to be overlapped between neuron samples and progenitor samples, we identify the caSNP-caPeak as an overlapped caQTL between cell types.

In a similar analysis for fetal brain eQTLs, we listed all SNPs with pairwise LD r^2^ > 0.5 with index eSNPs using LD matrices from fetal brain samples genotype data. Then we listed all SNPs with pairwise LD r^2^ > 0.5 with index caSNPs using LD matrices from caQTL samples genotype data. If any listed SNP is overlapped, we identify the eSNP-eGene and caSNP-caPeak as an overlapped QTL.

### Comparison to adult dorsolateral prefrontal cortex caQTLs

ATAC-seq peaks and caQTLs of adult dorsolateral prefrontal cortex (DLPFC) tissues were acquired from a previous study (Bryois et al., 2018). We used the same method from the section “Enrichment of peaks within annotated regions of the genome” above to calculate the enrichment of ATAC-seq peaks from this adult dataset at regulatory elements from different tissues. In addition to in vivo tissues/cell types, we also include cultured cells from (http://www.broadinstitute.org/~jernst/MODEL_IMPUTED12MARKS/). We found ATAC-seq peaks from cultured neural cells (neurons and progenitors) and DLPFC are enriched in regulatory elements of brain at different development stages, as expected. We found ATAC-seq peaks from cultured neural cells are highly enriched in enhancers and promoters of fetal brain tissues, but ATAC-seq peaks from DLPFC are highly enriched in enhancers and promoters of brain dorsolateral prefrontal cortex (Supplemental Figure 6A), even though 52% of ATAC-seq peaks from adult prefrontal cortex are overlapped with 39% of peaks from cultured neural cells (Supplemental Figure 6B). To calculate the overlap of caQTLs between cultured neural cells and DLPFC, we only kept bi-allelic SNPs and clumped neuron and progenitor caQTLs 5kb up-stream and 5kb down-tream from ATAC-seq peak centers as was previously done for caQTLs from DLPFC (Bryois et al., 2018). We identified overlapped caQTLs with the following process: 1) we identified ATAC-seq peaks (caPeaks) overlapped (by at least 1 bp) between the two datasets; 2) we selected the SNPs significantly associated with the same overlapped caPeak in both datasets; 3) we retained only LD-independent SNPs by: a) finding neuron/progenitor index caSNP-peak pairs (clumped) in the significant DLPFC caSNP-peak pairs (not-clumped), b) finding DLPFC caSNP-peak index caSNP-peak pairs (clumped) in the significant neuron/progenitor caSNP-peak pairs (not-clumped), c) we called an overlapped caQTL if either of the above (a) or b)) are true. We found highly temporal specificity in caQTLs (Supplemental Figure 6C).

### Determining the impact of caSNPs on motifs

In order to determine if genetic variation within peaks impacts transcription factor (TF) binding motifs, we used motifBreakR (v1.14.0) (Coetzee et al., 2015) to map known TF motifs to the sequence surrounding the neuron-specific/progenitor-specific significant caSNPs located in a ATAC-peak have significant association with the caPeak (parameter setting: threshold = 1e-4). All annotated motifs (in total 1304 TFs, 2943 motifs) are from MotifDb (1.26.0) (Shannon and Richards, 2014). We calculated the relative entropy (parameter setting: method=”ic”) for reference allele and alternative allele, then only keep the TFBSs which are strongly affected by the SNPs (motifbreakR parameter setting: effect=”strong”).

We calculated the ratios for the number of motif-disrupting SNPs that were neuron-specific significant/progenitor-specific significant bi-allelic caSNPs relative to the number of motif-disrupting bi-allelic SNPs that were located in accessible peaks for each TF regardless of whether there was a caQTL (Figure 6B).

To determine if the motif disrupting allele is associated with increased/decreased chromatin accessibility, we first identified the motif-disrupting allele. The motif-disrupting allele will decrease the relative entropy of the position possibility matrix of a TFBS. Then, we aligned the motif-disrupting allele with the effect allele for caQTLs. Finally, we used linear regression to determine the relationship between decreased relative entropy and effect size for all motif-disrupting alleles for this TFBS (lm(effect size ∼ decreased relative entropy+0). We fit the line through zero because we assume that if a motif is not disrupted by an allele, it will also have no effect on chromatin accessibility. The significance of the coefficient for effect size on decreased relative entropy was tested and the p-values adjusted to control FDR (Benjamini and Hochberg, 1995) (Figures 6C-F).

### Effect size correlation of eQTLs across different tissues

To compare effect size correlation between caQTLs and eQTLs, we calculated effect size correlation of eQTLs between brain cortex and other different tissues. eQTLs from different tissues (N=48, including brain cortex) were downloaded from GTEx (https://gtexportal.org/home/datasets). Significant eQTLs for all tissues were from GTEx version 7. We also downloaded tissue-specific “all SNP-Gene associations” for brain cortex and liver from GTEx version 7. To keep the most representative SNP-Gene pairs, we only retained the most significant SNP for each eGene in every tissue as the significant eQTLs.

We found the same SNP-Gene pairs from brain cortex with the significant eQTLs from liver, and calculated effect size (“Slope” in the files) correlation for these eQTLs, and vice versa for liver eQTLs (Figure 5F). Then, we calculated effect size correlation between the same SNP-Gene pairs from brain cortex with the significant eQTLs from other tissues (N=45), except Brain_Cortex, Brain_Frontal_Cortex_BA9 and Brain_Anterior_cingulate_cortex_BA24 (Figure 5G).

For caQTLs, we only retained the most significant caSNP for each caPeak in progenitors and neurons, then calculated effect size correlation between the same SNP-Peak pairs in neurons with the significant caQTLs in progenitors, and vice versa for significant caQTLs in neurons (Figure 5E).

### Partitioned Heritability

Partitioned heritability was measured using LD Score Regression v1.0.0 (Finucane et al., 2015b) to identify enrichment of GWAS summary statistics among differentially accessible peaks. First, an annotation file was created which marked all HapMap3 SNPs that fell within Neuron>Progenitor or Progenitor>Neuron differentially accessible peaks. To avoid the bias caused by different numbers of progenitor > neuron and neuron > progenitor differentially accessible peaks, we randomly selected the same number of Progenitor>Neuron as were significant Neuron>Progenitor peaks. LD-scores were calculated for these SNPs within 1 cM windows using the 1000 Genomes data. These LD-scores were included simultaneously with the baseline distributed annotation file from (Finucane et al., 2015b). Subsequently, the heritability explained by these annotated regions of the genome was assessed from these genome-wide association studies: Attention-Deficit/hyperactivity disorder (Demontis et al., 2019), autism spectrum disorder (Grove et al., 2019), IQ (Savage et al., 2018), major depressive disorder (Wray et al., 2018), Bipolar disorder (Stahl et al., 2019), schizophrenia (Pardiñas et al., 2018), insomnia (Jansen et al., 2019b), educational attainment (Lee et al., 2018a), subjective well-being (Okbay et al., 2016), depressive symptoms (Okbay et al., 2016), neuroticism (Nagel et al., 2018), anorexia nervosa (Duncan et al., 2017), anxiety (Otowa et al., 2016), Alzheimer’s disease (Jansen et al., 2019a), epilepsies (International League Against Epilepsy Consortium on Complex Epilepsies, 2018), Parkinson’s disease (Nalls et al., 2018).

The enrichment was calculated as the heritability explained for each phenotype within a given annotation divided by the proportion of SNPs in the genome and Benjamini–Hochberg FDR correction (Benjamini and Hochberg, 1995) was used to correct for multiple comparisons.

### Co-localization with GWAS data

We used conditional caQTLs to detect the co-localization of caQTLs and multiple GWAS data as previously listed above in **Partitioned Heritability**. First, to identify co-localized loci: 1) we listed SNPs with pairwise LD r^2^ > 0.8 with the caSNPs in the caPeak using genotype data from neuron samples and progenitor samples, separately; 2) we listed SNPs with pairwise LD r^2^ > 0.8 with index GWAS SNP (p<5e-8 and exhibited the strongest association in upstream/downstream 100kb from the center of this caPeak) using the LD matrix from European genotype data from 1000 Genome project phase 3 with population code EUR. Second, we labelled the caPeak as a potentially co-localized loci if any SNP from the above two categories is overlapped. Third, we performed conditional caQTL for significant (FDR < 0.05) caSNPs in the potential co-localized locus (caPeak) conditioned on the index GWAS SNP. If the caQTL is no longer significant (FDR > 0.05), then we identified the caPeak as a co-localized loci with this GWAS trait.

